# Development of an HIV reporter virus that identifies latently infected CD4^+^ T cells

**DOI:** 10.1101/2022.01.17.476679

**Authors:** Eun Hye Kim, Lara Manganaro, Michael Schotsaert, Brian D. Brown, Lubbertus C.F. Mulder, Viviana Simon

**Affiliations:** Department of Microbiology, Icahn School of Medicine at Mount Sinai, New York, USA, 10029; Division of Infectious Disease, Department of Medicine, Icahn School of Medicine at Mount Sinai, New York, USA; The Global Health and Emerging Pathogens Institute, Icahn School of Medicine at Mount Sinai, New York, USA; INGM-Istituto Nazionale Genetica Molecolare, Milan, Italy, 20122; Department of Pharmacological and Biomolecular Sciences (DiSFeB), University of Milan, Milan, Italy; Department of Pathology, Molecular and Cellular Medicine at Icahn School of Medicine at Mount Sinai, New York, USA, 10029; Precision Immunology Institute, Icahn School of Medicine at Mount Sinai, New York, USA, 10029; Department of Genetics and Genomic Sciences, Icahn School of Medicine at Mount Sinai, New York, 10029

**Keywords:** HIV-1, Latency establishment, CD4^+^ T cell subsets, Reporter virus, HIV integration, Mass cytometry

## Abstract

There is no cure for HIV infection as the virus establishes a latent reservoir, which escapes highly active antiretroviral treatments. One major obstacle is the difficulty to identify cells that harbor latent proviruses. We devised a novel viral vector that carries a series of versatile reporter molecules that are expressed in an LTR-dependent or LTR-independent manner and allows to accurately distinguish productive from latent infection. Using primary human CD4^+^ T cells, we show that transcriptionally silent proviruses make up over 50% of all infected cells. These latently infected cells harbor proviruses, but lack evidence for viral transcription. LTR silent integrations occurred to variable degrees in all CD4^+^ T-subsets examined with CD4^+^ T_EM_ and CD4^+^ T_reg_ displaying the highest frequency. Viral vectors such as the one described here, permit interrogation HIV latency at a single-cell resolution, revealing mechanisms of latency establishment and allowing for the characterization of effective latency reversing agents.

## Introduction

An HIV/AIDS cure remains elusive because HIV establishes a hard to eradicate latent reservoir. Indeed, despite the intense efforts of the scientific community over the past forty years, one of the main obstacles to HIV eradication is the difficulty to identify the cells that harbor transcriptionally silent proviruses forming a long-lasting, antiretroviral resistant, latent reservoir.

CD4^+^ T cells are the primary targets of HIV infection and contribute to establishing and maintaining the latent reservoir (Churchill et al., 2016). Subsets of these cells differ in their susceptibility to infection, their ability to traffic, exert effector functions, proliferate, and survive for prolonged periods of time (Zerbato et al., 2016, Kulpa et al., 2019, Grau-Exposito et al., 2019). A more detailed and cell-type specific understanding of latency establishment is, therefore, of great relevance.

HIV latency can result from progressive epigenetic silencing of productive infections (Colin and Van Lint, 2009, Siliciano and Greene, 2011, Pearson et al., 2008, Weinberger et al., 2008, Hashemi et al., 2016, Jefferys et al., 2021, Mbonye and Karn, 2014). However, evidence suggest that a substantial portion of HIV integrations occur in an LTR-silent manner (Dahabieh et al., 2013, Chavez et al., 2015, Calvanese et al., 2013). This shift in our understanding of HIV latency establishment was made possible by the use of HIV vector constructs, that encode fluorescent reporters whose expression are either dependent or independent of HIV LTR promoter activity (Dahabieh et al., 2013, Calvanese et al., 2013). We reported previously that LTR from different HIV subtypes differ in their propensity for silencing (Dahabieh et al., 2013) and that latent infection was inversely correlated with cellular activation and NF-κB activity (Dahabieh et al., 2014), but these studies were limited to immortalized T cell lines.

A number of different dual-reporter vectors have been used in the past to interrogate different aspects of latency establishment (Battivelli et al., 2018, Calvanese et al., 2013, Cavrois et al., 2017, Chavez et al., 2015, Dahabieh et al., 2014, Dahabieh et al., 2013, Hashemi et al., 2016, Kim et al., 2019, Matsuda et al., 2015, Cai et al., 2021). They all rely on the LTR-dependent or LTR-independent differential expression of reporter proteins. We reasoned that positioning the reporters in distinct reading frames undergoing different splicing events would reduce the potential for promoter interference. Moreover, we decided that having full-length functional *gag* and accessory proteins, including *Nef*, would be necessary to obtain a physiologically relevant assessment of HIV latency establishment. Lastly, we chose the strong, constitutively active PGK, promoter to ensure high expression of the latency protein tags (Jones et al., 2009, Kim et al., 2019, Cai et al., 2021). Of note, the resulting HIV molecular clone, pMorpheus-V5, lacks a functional envelope, which eases biosafety precautions for down-stream applications with live cells and allows for pseudotyping with different HIV envelopes.

Detection of infected cells at the single cell-level in a manner that is independent of HIV LTR activity allows probing for potential differences in latency establishment in specific CD4^+^ T cell subsets. We validated our system by analyzing primary human CD4^+^ T cells, stimulated with IL-2 and *α−*CD3/CD28 or IL-2 alone, infected with pMorpehus-V5 by both conventional flow cytometry as well as high-dimensional cytometry by time-of flight (CyTOF). Productively infected cells expressed the four encoded reporters with equal efficiency, suggesting that promoter interference was not an issue with this HIV reporter vector. Furthermore, down-regulation of the CD4 receptor by HIV *Nef* was only observed in cells co-expressing all the four tags while cells expressing only the PGK driven protein tags displayed CD4 receptor levels comparable to uninfected cells.

In summary, we show here that primary human CD4^+^ T cells harboring transcriptionally silent proviruses are more frequent than productively infected cells. This was true for all seven CD4^+^ T cell subsets investigated in this study but CD4^+^ regulatory T cells (CD4^+^ T_REG_) and CD4^+^ effector memory T cells (CD4^+^ T_EM_) had the highest frequency of LTR silent infection. We propose that the pMorpheus-V5 system encoding all the viral accessory proteins as well as different reporter proteins will facilitate obtaining new mechanistic insights into latency establishment and reactivation in different primary human cells including but not limited to primary human CD4^+^ T cells.

## Results

### pMorpheus-V5 construction and identification of productively and latently infected cells

To investigate the cellular processes that govern establishment of HIV latency at the single-cell level in human primary CD4^+^ T cells, we developed a viral reporter vector (pMorpheus-V5) suitable to detect the LTR dependent or independent expression of four reporters, two of which are membrane bound (**Figure 1A**). A PGK promoter driven reporter cassette encoding V5-NGFR was inserted into *Env* open reading frame effectively deleting a large portion of the HIV envelope. Upon proviral integration, but independently of LTR promoter activation, V5-NGFR localizes to the surface of the infected cell. The LTR-dependent expression cassette includes sequences encoding HSA and mCherry that were cloned upstream of an internal ribosomal site followed by *Nef*. Thus, LTR-dependent expression of HSA, mCherry and *Nef* are linked (Young et al., 2018). Cells productively infected with pMorpheus-V5 express four reporters: V5-NGFR driven by PGK as well as HSA and mCherry driven by the HIV-LTR. In contrast, latently infected cells express only V5-NGFR. The combination of fluorescence reporter and membrane bound tags allow for easy and specific identification of the different cell population by flow cytometry (**Figure 1B, 1C**). Cells expressing only HSA and mCherry (lower right quadrant in the cartoon) should not be found since all productively infected cells harbor proviruses and, thus, would be positive for V5-NGFR (**Figure 1B**). To assess the ability to reliably distinguish cells harboring LTR-silent or LTR-active proviruses, we infected primary human CD4^+^ T cells, stimulated either with IL-2 alone or activated by a combination of IL-2 and *α*-CD3/CD28 coated beads, with pMorpheus-V5. We produced pMorpheus-V5 viral stocks by co-transfecting the molecular clone with a dual-tropic envelope pSV III-92 HT593.1 (Gao et al., 1996). Primary human CD4^+^ T cells were infected with pMorpheus-V5 and followed for five days. We used flow cytometry or CyTOF for the analysis. The cell viability was comparable between infected and uninfected cells under each experimental condition (**Figure S1A, S1B, Figure S2A, S2B**). NGFR and V5 as well as HSA and mCherry provided very comparable percentages of latently or productively infected cells as measured by either flow-or mass-spectrometry (**Figure 1C**: Flow cytometry, **Figure S2C**: CyTOF). The absence of HSA/mCherry expressing cells lacking V5-NGFR expression indicates that promoter interference is not a problem with the current vector system.

**Figure 1.**
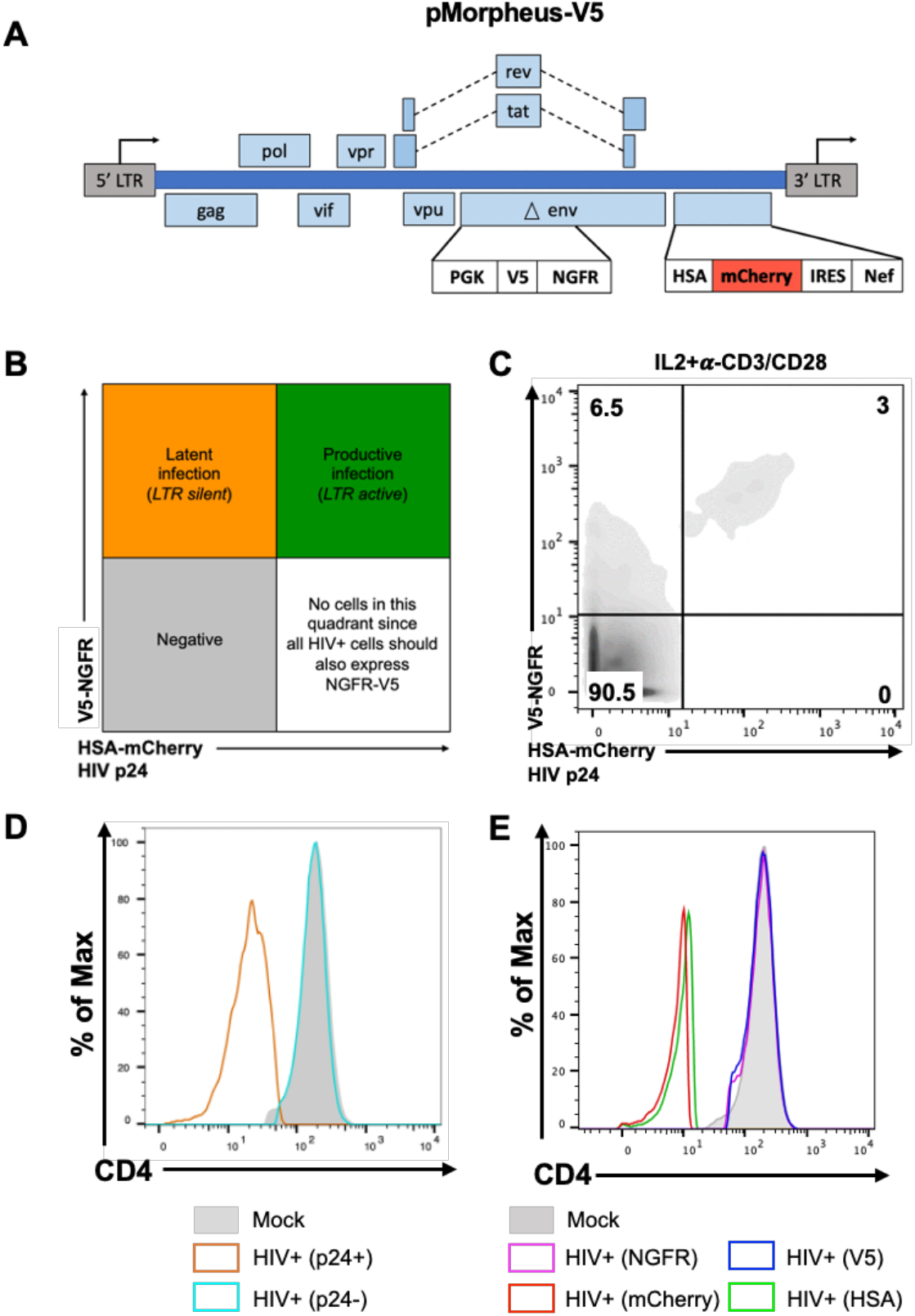
Description and characterization of the pMorpheus-V5 viral reporter vector. (A) Schematic representation of the HIV reporter viral vector pMorpheus-V5 (pLAI2-V5-NGFR HSA-mCherry-IRES-*Nef*). (B) The gating strategy to identify cells latently or productively infected by pMorpheus-V5 by flow cytometry or CyTOF is depicted. (C) Latently (V5+, NGFR+) or productively (HSA_POS_, mCherry_POS_, V5_POS_, NGFR_POS_, p24_POS_) infected, IL-2+*α*-CD3/CD28 stimulated CD4^+^ T cells were identified by flow cytometry five days after infection with pMorpheus-V5. The data shown is one representative example of four independent experiments using cells from different healthy donors. The percentage of CD4^+^ T cells in each quadrant are indicated. Representative data from four independent donors are shown. (D) *Nef* down-regulates CD4 levels during pMorpheus-V5 infection. CD4^+^ T cells from two different donors were stimulated with IL-2+*α*-CD3/CD28 and infected with pMorpheus-V5. CyTOF was performed 5 days post-infection. Histogram plots show the CD4 expression for uninfected cells (shaded light grey), p24_POS_ cells (orange), p24_NEG_ cells (light blue). The data were analyzed with FlowJo. These results are representative of data from two independent donors. (E) Cells were stimulated with IL-2+*α*-CD3/CD28 prior to infection with p-Morpheus-V5. Histogram analysis of CD4 expression levels of NGFR (magenta), V5 (blue), mCherry (red), or HSA (green) positive cells. The data were analyzed by FlowJo. Representative data from two independent donors are shown.

Since HIV-1 *Nef* is known to down-regulate the CD4 receptor (Garcia and Miller, 1991, Guy et al., 1987), we next compared the expression of CD4 in latently and productively infected CD4^+^ T cells with that of mock infected cells. As expected, HIV Gag p24 positive CD4^+^ T cells expressed substantially less CD4 compared to p24 negative cells or mock infected cells (**Figure 1D, 1E** (CyTOF), and **Figure S1C, 1D** (Flow cytometry), **Figure S2D, 2E** (CyTOF)). Down-regulation of CD4 was also observed in HSA_POS_ or mCherry_POS_ cells but not in NGFR_POS_ or V5_POS_ only cells. Thus, CD4 expression in CD4^+^ T cells harboring only LTR-silent proviruses (as identified by NGFR_POS_/V5_POS_ but HSA_NEG_/mCherry_NEG_ expression) remained comparable to that found on mock-infected cells (**Figure 1D, 1E** and **Figure S2D, 2E** (CyTOF), **Figure S1C, 1D** (Flow cytometry)).

Taken together, infection with pMorpheus-V5 permits to distinguish between latently infected and productively infected primary CD4^+^ T cells using either flow cytometry or CyTOF.

### Primary human CD4^+^ T cells harboring LTR silent proviruses are as frequent as those harboring LTR active proviruses

We next measured the proportion of latently and productively infected CD4^+^ T cells five days after infection with pMorpheus-V5. We specifically compared the expression of the different viral reporters and/or proteins including mCherry and/or intracellular p24, NGFR and V5 in CD4^+^ T cells obtained from four independent healthy donors (**Figures 2A** and **2B**). The proportion of cells harboring LTR-silent proviruses was significantly higher than the number of productively infected in IL-2-only stimulated cells (**Figure 2C**, average LTR-silent 58%±4 versus average LTR-active 42%±4, *p*=0.0014). This difference was reduced when CD4^+^ T cells were activated with IL-2±*α*-CD3/CD28 (**Figure 2D**, average LTR-silent 52%±8 vs average LTR-active 48%±8, *p* =0.61).

Thus, more than half of the infected primary human CD4^+^ T cells harbor LTR-silent proviruses in a manner that is independent of the type of stimulation used with very limited inter donor variation (**Figure 2C, 2D**).

**Figure 2.**
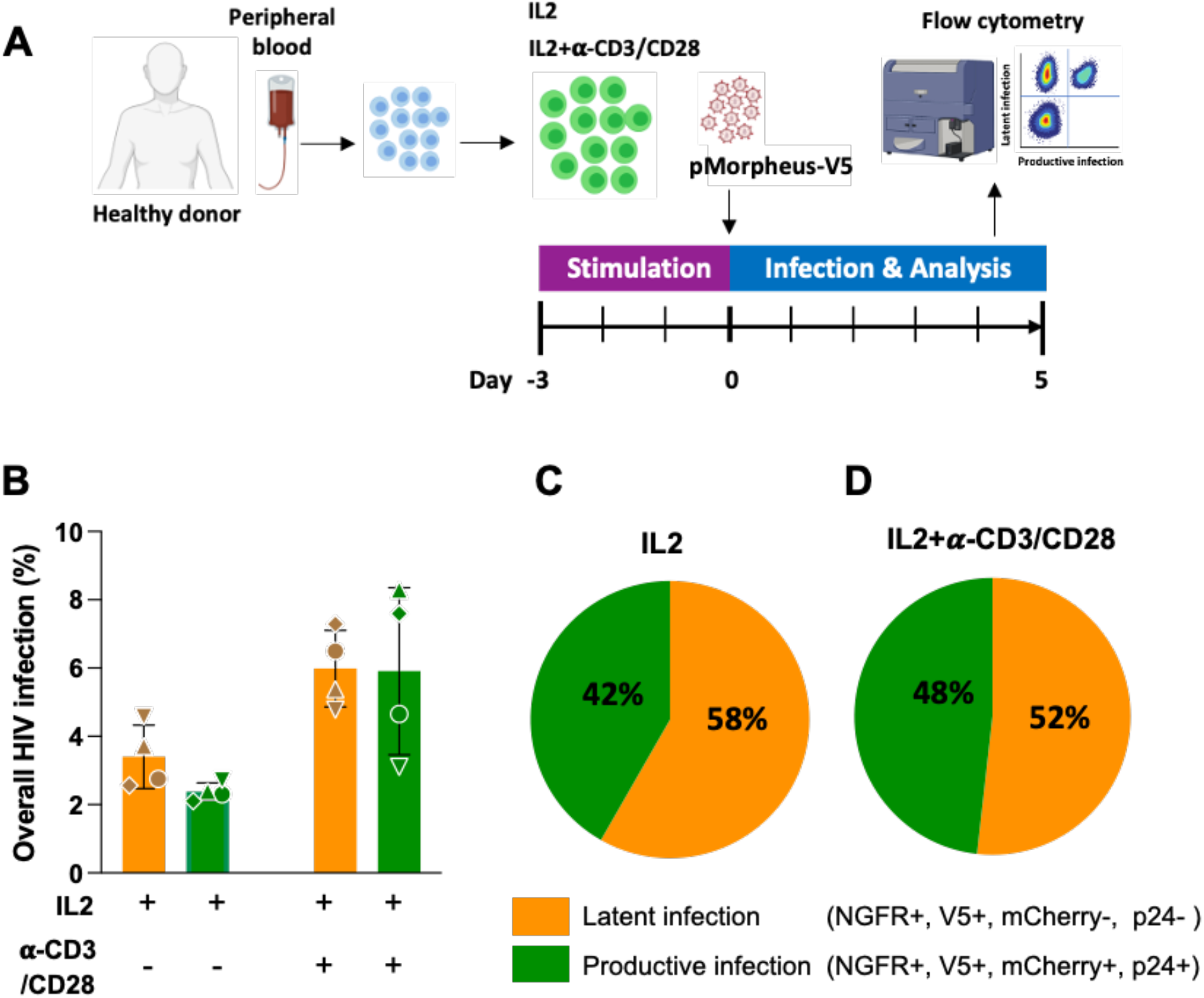
Identification of latently and productively infected CD4^+^ T-cells. (A) Schematic representation of the experimental design to analyze CD4^+^ T cells infected with pMorpheus-V5. CD4^+^ T cells from four healthy donors were stimulated with IL-2 or a combination of IL-2 with *α*-CD3/CD28 antibody-coated beads for 3 days prior to infection with pMorpheus-V5. The frequency of latently and productively infected cells was determined five days post-infection by Flow Cytometry. (B) The frequency of latently (V5_POS_, NGFR_POS_, yellow) and productively (mCherry_POS_, V5_POS_, NGFR_POS_, p24_POS_, green) infected CD4^+^ T cells stimulated with IL-2 alone or IL-2/*α*-CD3/CD28 is shown. Cells were analyzed by flow cytometry five days after infection. Each of the four healthy donors is identified by symbol. Error bars represent the average ±SD. (C) The relative proportion of pMorpheus-V5 latently (yellow) or productively (green) infected CD4^+^ T cells stimulated with IL-2 is shown. The data represent the average of the four individual healthy donors. (D) The relative proportion of pMorpheus-V5 latently (yellow) or productively (green) infected CD4^+^ T cells activated with *α*-CD3/CD28 is shown. The data represent the average of the four individual healthy donors.

### Detection of HIV integration and transcription in latently infected primary human CD4^+^ T cells

We next measured the level of integration and viral transcription in LTR-silent and LTR-active infected CD4^+^ T cells (**Figure 3A**). Briefly, we infected IL-2+*α*-CD3/CD28 stimulated primary human CD4^+^ T cells from three different healthy donors with pMorpheus-V5 in the presence and absence of the FDA-approved integrase inhibitor Raltegravir. Five days later, culture supernatants were collected for p24 analysis and the live CD4^+^ T cells were sorted by FACS (latently infected: V5-NGFR_POS_/HSA-mCherry_NEG_; productively infected: V5-NGFR_POS_/HSA-mCherry_POS_, as well as cells lacking reporter expression: NGFR_NEG_/HSA-mCherry_NEG_). Raltegravir prevented both latent as well as productive infections (**Figure 3B**).

**Figure 3.**
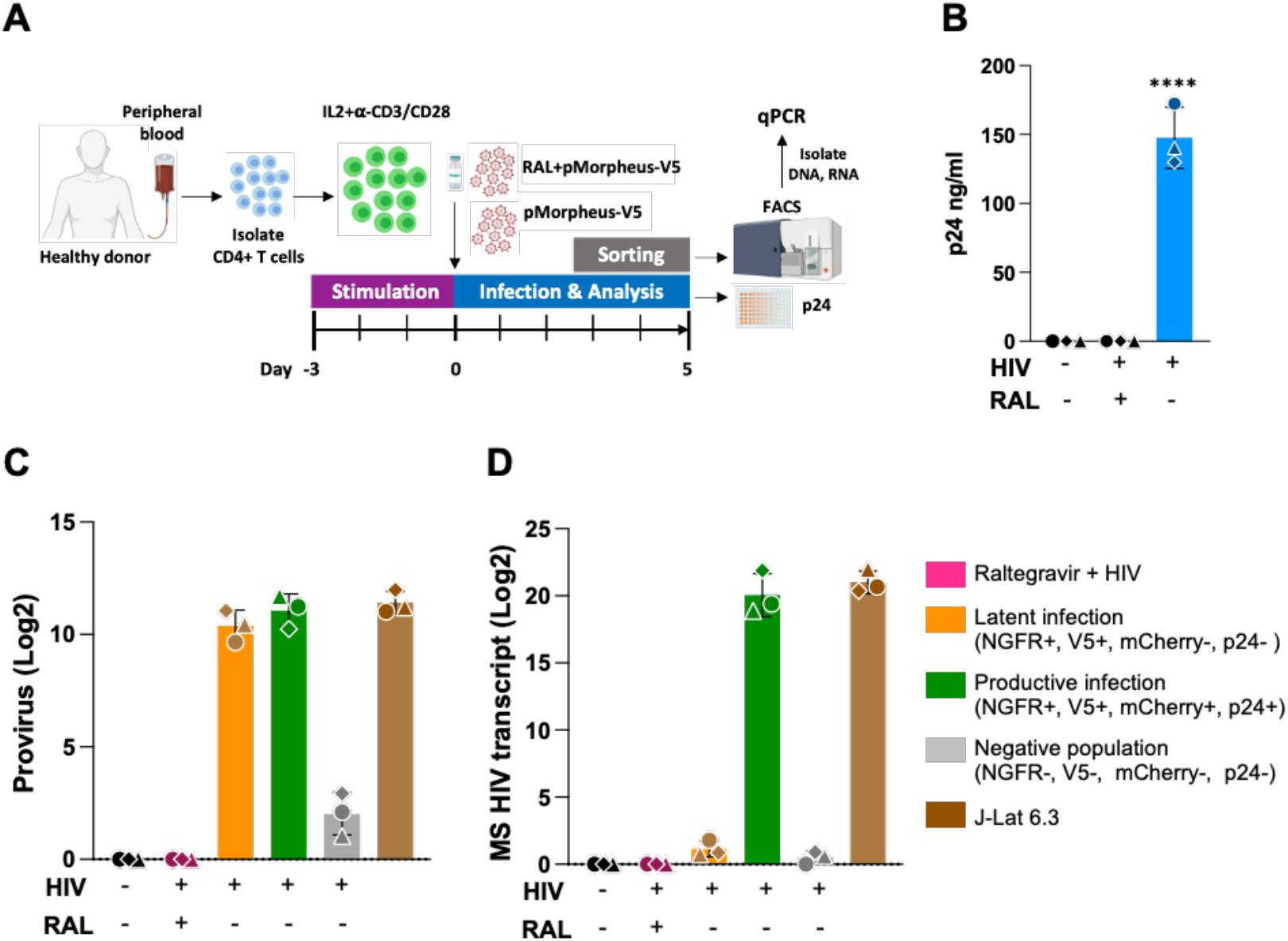
Quantification of proviruses and multiply spliced HIV transcripts in FACS-sorted CD4^+^ T cells. (A) Schematic representation of the experimental design to analyze provirus integration or multiply spliced transcripts in productive or latently infected cells from three different healthy donors. CD4^+^ T cells were activated with IL-2 and *α*-CD3/CD28 antibody-coated beads for 3 days prior to infection with pMorpheus-V5. CD4^+^ T cells were infected with pMorpheus-V5 in the presence and absence of Raltegravir. Latently (orange), productively (green) and the reporter negative cell populations were sorted by FACS five days post-infection with pMorpheus-V5. RNA and DNA was extracted from the same samples. (B) Gag p24 concentration was measured by ELISA in the culture supernatants five days after infection with pMorpheus-V5. Bar graph shows the production of p24 in pMorpheus-V5 or mock infected cells in the presence and absence of Raltegravir (RAL). Results for each of the three healthy donors are identified by a specific symbol. The average of three donors is shown (±SD), **** denotes *p*<0.0001. (C) HIV integration was measured by a two-step quantitative *Alu*-PCR assay using 100ng of total cellular DNA from each sorted cell population (Latent infection: yellow, productive infection: green; reporter expression negative cells: grey). The bar graph depicts relative amounts of integrated proviruses. DNA from J-Lat 6.3 was used as a positive control. The mean ±SD of three independent experiments/donors are shown. Sorted cells from the same healthy donor (N=3) are identified by the same symbol. D) Multiply spliced (MS) HIV transcripts were quantified by semi-nested real-time quantitative PCR. 100 ng of RNA was extracted from the same samples as in (C) to synthesize cDNA. RNA extracted from stimulated J-Lat 6.3 was used as a positive control. The bar graph depicts the means ±SD from three independent experiments/donors. Sorted cells from the same healthy donor (N=3) are identified by the same symbol.

Genomic DNA was extracted from sorted cells to measure integration using *Alu-*1 PCR (Butler et al., 2001), while cellular RNA was extracted from the same samples to detect multiply spliced *tat/rev* HIV transcripts by RT-qPCR assay (Pasternak et al., 2008, Zerbato et al., 2021) (**Figure 3A**). As a positive control, we used the T cell-line J-Lat, which harbors a single provirus, stimulated with phorbol 12-myristate 13-acetate (PMA, 16 nM,) and Ionomycin (0.5 μM) for 3 days. We found that the levels of integrated provirus were comparable between latently and productively infected (**Figure 3B**). As expected, Raltegravir treatment effectively blocked HIV integration. Interestingly, a small portion of the cells lacking expression of all four reporters (“negative cell population”) do carry integrated proviruses pointing to the fact that the pMorpheus-V5 vector system may still miss a small number of integration events possibly by less than intact proviruses.

To assess the level of HIV transcription, we measured multiply spliced HIV transcripts in each of the sorted CD4^+^ T cell populations (with and without Raltegravir treatment). As shown in **Figure 3D**, multiply spliced HIV transcripts were only detected in the productively infected CD4^+^ T cells (V5-NGFR_POS_/HSA and mCherry_Pos_) and not in the LTR-silent (V5-NGFR_POS_) or reporter negative CD4^+^ T cells. Similarly, the Raltegravir treated cells lacked expression of multiply spliced HIV transcripts while the stimulated J-Lat cells expressed multiply spliced HIV transcripts to levels comparable to those found in productively infected primary human CD4^+^ T cells.

Collectively, these data reinforce the notion that the difference between LTR-active and LTR-silent cells are solely based on transcription and not on reduced levels of viral integration.

### Identification of CD4^+^ T cell subpopulations that preferentially support HIV latency

Since pMorpheus-V5 allows to distinguish between LTR-silent and LTR-active infection events, we next used cytometry by time-of-flight (CyTOF) to determine whether CD4^+^ T cell subpopulations differ in their propensity to support latency establishment. Primary human CD4^+^ T cells stimulated with IL-2 alone or IL-2-*α*-CD3/CD28 were infected with pMorpheus-V5 and stained for CyTOF five days later. We first barcoded the individual infections with CD45 antibodies, and then stained with a custom antibody cocktail recognizing the HIV-encoded reporters as well as 29 other cellular markers (**Figure S3A**). Multi-dimensional analysis identified the protein tags of latent infection (V5-NGFR_POS_ only) in more 50% of the cells analyzed (**Figures 4A, 4C, 4D, 5A and 5B**). The relative percentage of productively infected cells increased overall with activation (compare IL2-*α*-CD3/CD28 to IL2 alone in **Figure 4B**) while the proportion of latently infected cells was higher in the IL-2-only stimulated cells compared to the IL-2-*α*-CD3/CD28 stimulated cells (**Figures 4C and 4D**). We visualized the latently and productively infected cells at the single-cell level using viSNE, which employs *t*-stochastic neighbor embedding (*t*-SNE) to generate a two-dimensional map of the latency protein tag expression in different CD4^+^ T cell populations (**Figure 4E**).

**Figure 4.**
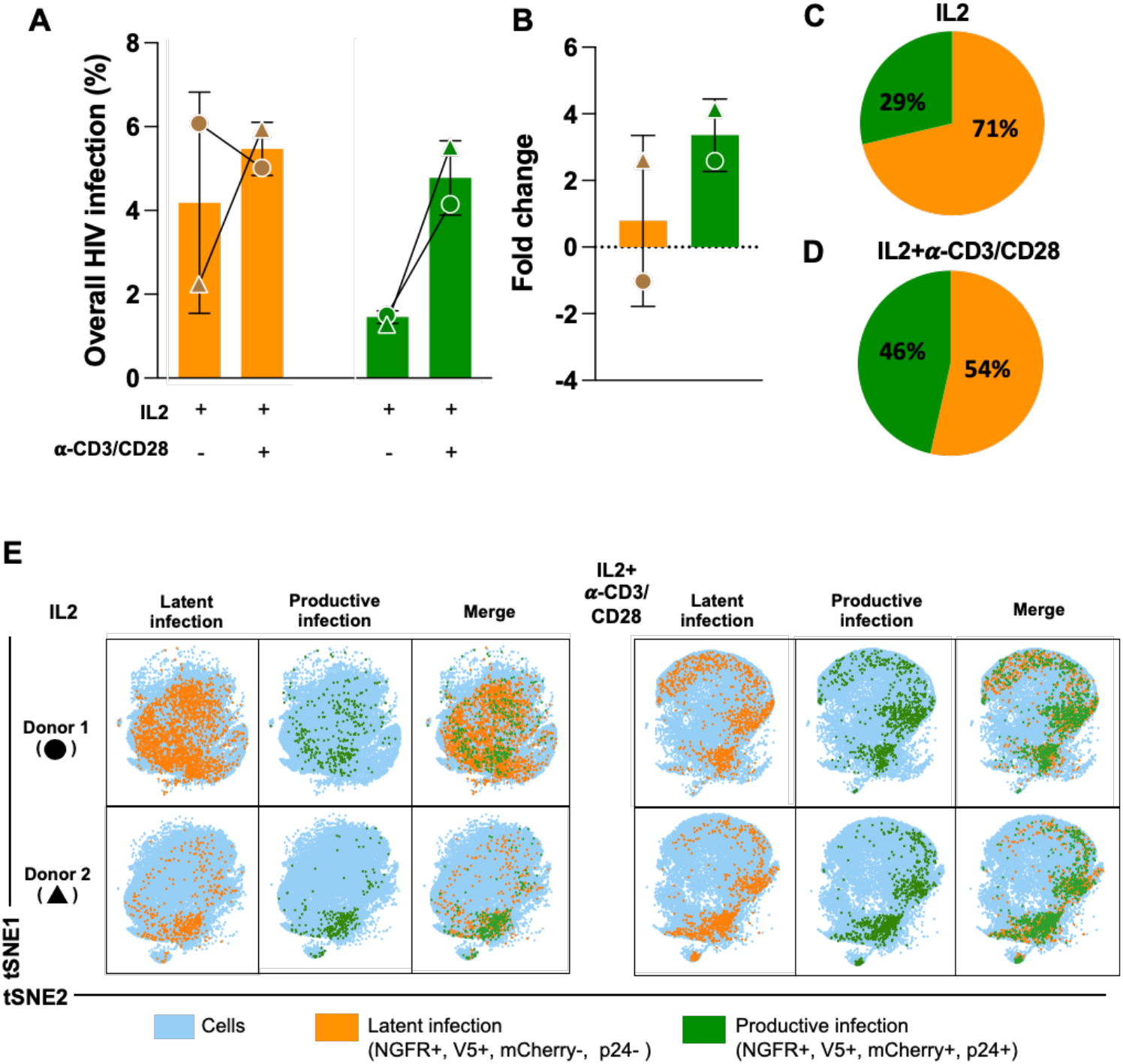
Characterization of latently and productively infected CD4^+^ T cell populations using Mass Cytometry. (A) pMorpheus-V5-infected primary human CD4^+^ T cells stimulated with IL-2 or activated with IL-2/*α*-CD3/CD28 beads were analyzed by mass cytometry five days after infection. Latently (V5_POS_, NGFR_POS_, yellow) or productively (HSA_POS_, mCherry_Pos_, V5_POS_, NGFR_POS_, p24_POS_, green) infected cells identified. Data for each of the two healthy donors (±SD) is indicated by a symbol. (B) The bar graph shows the fold change of latently or productively infected cells depending on cellular activation. The average of two individual donors is shown. (C) The proportion of latently or productively infected CD4^+^ T cells stimulated with IL-2 for three days. The average of two individual donors is shown with the percentage and the total number of cells. (D) The proportion of latently or productively infected CD4^+^ T cells stimulated with IL-2 and *α*-CD3/CD28 beads for three days. The average of two individual donors is shown with the percentage and the total number of cells. (E) viSNE analysis of pMorpheus-V5 infected CD4^+^ T cells stimulated with IL-2 alone or with IL-2 and *α*-CD3/CD28. The tSNE1 and tSNE2 axes are based on the relevant markers of the subsets defined by the Cytobank program. The results are based on 66,000 cells for each condition from each donor identified by the circle and triangle symbols in panel A.

The CyTOF antibody panel used allowed identification of seven different CD4^+^ T cell populations. We noted that activation with IL2+*α*-CD3/CD28 modified the distribution of the CD4^+^ T cell subsets in the absence of pMorpheus-V5 infection (**Figure S4A, S4B, S4C**). We visualized the distribution of seven different CD4^+^ T cell subsets at single-cell level using viSNE, which employed *t*-stochastic neighbor embedding (*t*-SNE) to generate a two-dimensional map where the expression between cells corresponded to their marker profile (**Figure S4D and S4E**) in the presence of IL-2 or IL-2+*α*-CD3/CD28.

Across these subsets, the proportion of cells harboring LTR-silent proviruses (V5-NGFR_POS only_) were twice as high as the cells displaying productive infections (**Figure 5A and 5B**). We noted that LTR-silent infections were more common in cells stimulated with IL-2 while cell activation with IL-2+*α*-CD3/CD28 reduced the relative proportion of latently infected cells (**Figure 5B, 5C**). viSNE maps further visualize LTR-silent and LTR-active proportions in each distinct CD4^+^ T cell populations (**Figure S5A, S5B**). Of note, the vast majority of infected T_N_ are latent. We observed that CD4^+^ T_EM_ and CD4^+^ T_reg_ have the highest total number of latently infected cells and interestingly, IL-2/*α−*CD3/CD28 stimulation increases the percentage of productive infection in all subsets with exception of T_EM_. Taken together, our data indicate as similar finding in previous (Buzon et al., 2014) that latent infections occur across all seven CD4^+^ T cell subsets analyzed with T_EM_, T_REG_ contributing heavily to the latent reservoir formation.

**Figure 5.**
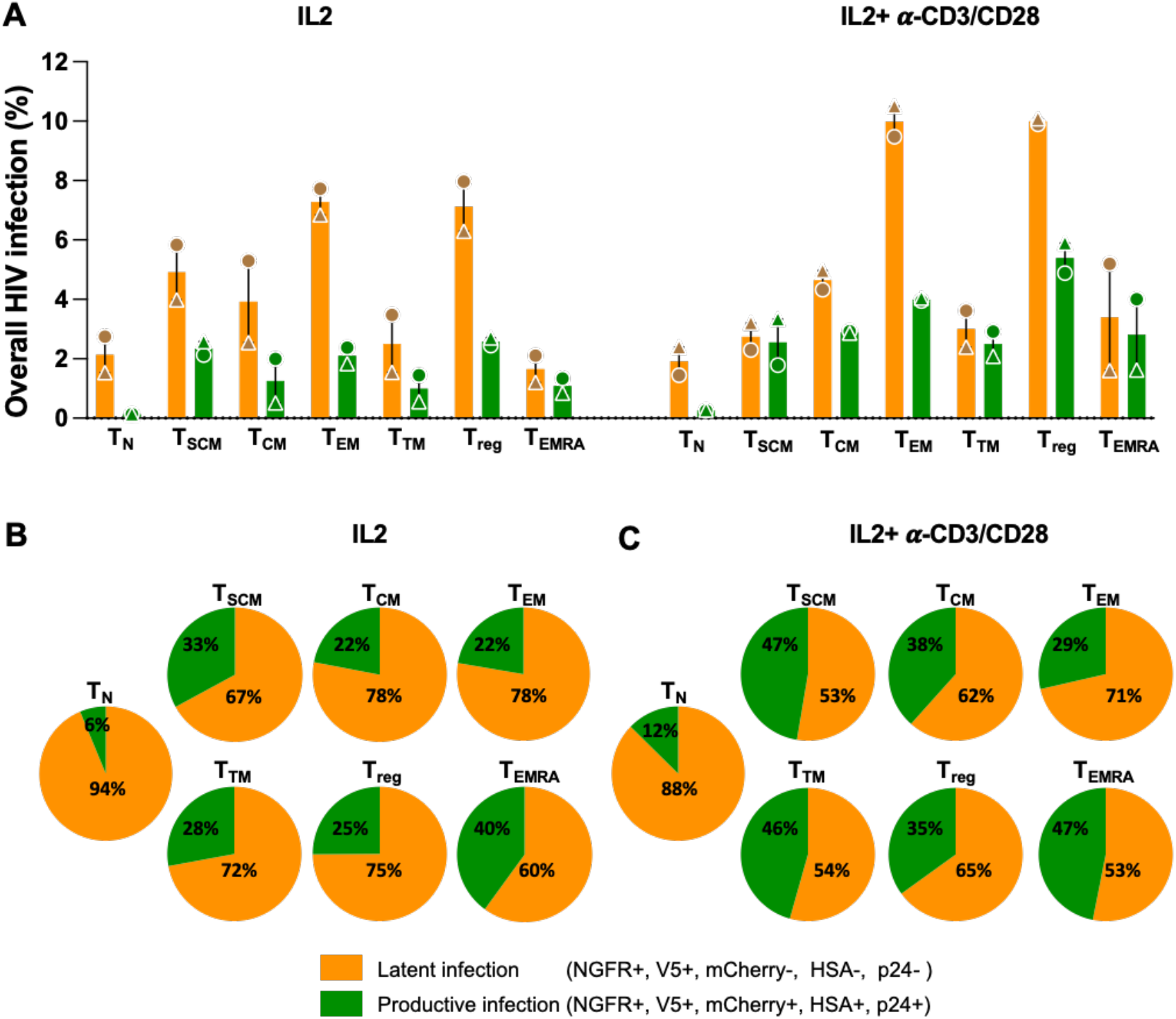
Determination of latent and productive infection in different CD4^+^ T cell subsets. (A) pMorpheus-V5-infected primary human CD4^+^ T cells from healthy donors stimulated with IL-2 or with IL-2 and *α*-CD3/CD28 antibody-coated beads were analyzed by CyTOF to determine levels of NGFR, V5, mCherry, HSA, and p24 in different CD4^+^ T cell subsets. The bar graphs represent percentage of latent or productive infection of each T cell subsets. The average of two individual donors is shown (±SD). Each donor is indicated by a unique symbol and each sorted group is color coded in the figure. The average of two individual donors is shown. (B) The proportion of latent (yellow) and productive (green) infection of CD4^+^ T cells upon pMorpheus-V5 infection. CD4^+^ T cells from 2 donors were stimulated with IL-2 for 3 days. The average of two individual donors is shown. (C) CD4^+^ T cells were stimulated with IL2 and *α*-CD3/CD28 beads for 3 days and infected with pMorpheus-V5. The proportion of latently (yellow) or productively (green) infected cells are depicted. The average of two individual donors is shown.

### Expression of activation markers confirms the latent and productive nature of infections with pMorpheus-V5

Expression of CD69 and HLA-DR associated with productive infection (Marini et al., 2008, Sahu et al., 2006) but not with latent infection (Moso et al., 2019). We therefore analyzed, the expression of the activation markers CD69 and HLA-DR in latently and productively infected CD4^+^ T cells upon pMorpheus-V5 infection. We observed that both markers were expressed at higher levels in productively infected CD4^+^ T cells compared to latently infected, the reporter negative cell populations or mock infected cells (**Figure 6A** and **6C**). In the absence of overt T cell activation, all the memory T cells populations showed higher expression of these two activation markers when productively infected (**Figure 6B**). The same was true following stimulation with IL-2+*α*-CD3/CD28 with the exception of CD4^+^ T_TM_ cells. This T cell subset displayed only a marginally higher expression of CD69/HLA-DR in productively infected cells compared to latently infected, the reporter negative populations or mock infected cells (**Figure 6D**). Importantly, latently infected CD4^+^ T cells showed the CD69/HLA-DR expression profiles that were comparable to the reporter negative cell populations or mock infected cells providing validation for the fact that latently infected cells identified by pMorpheus-V5 infection are fundamentally distinct from the productively infected cells.

**Figure 6.**
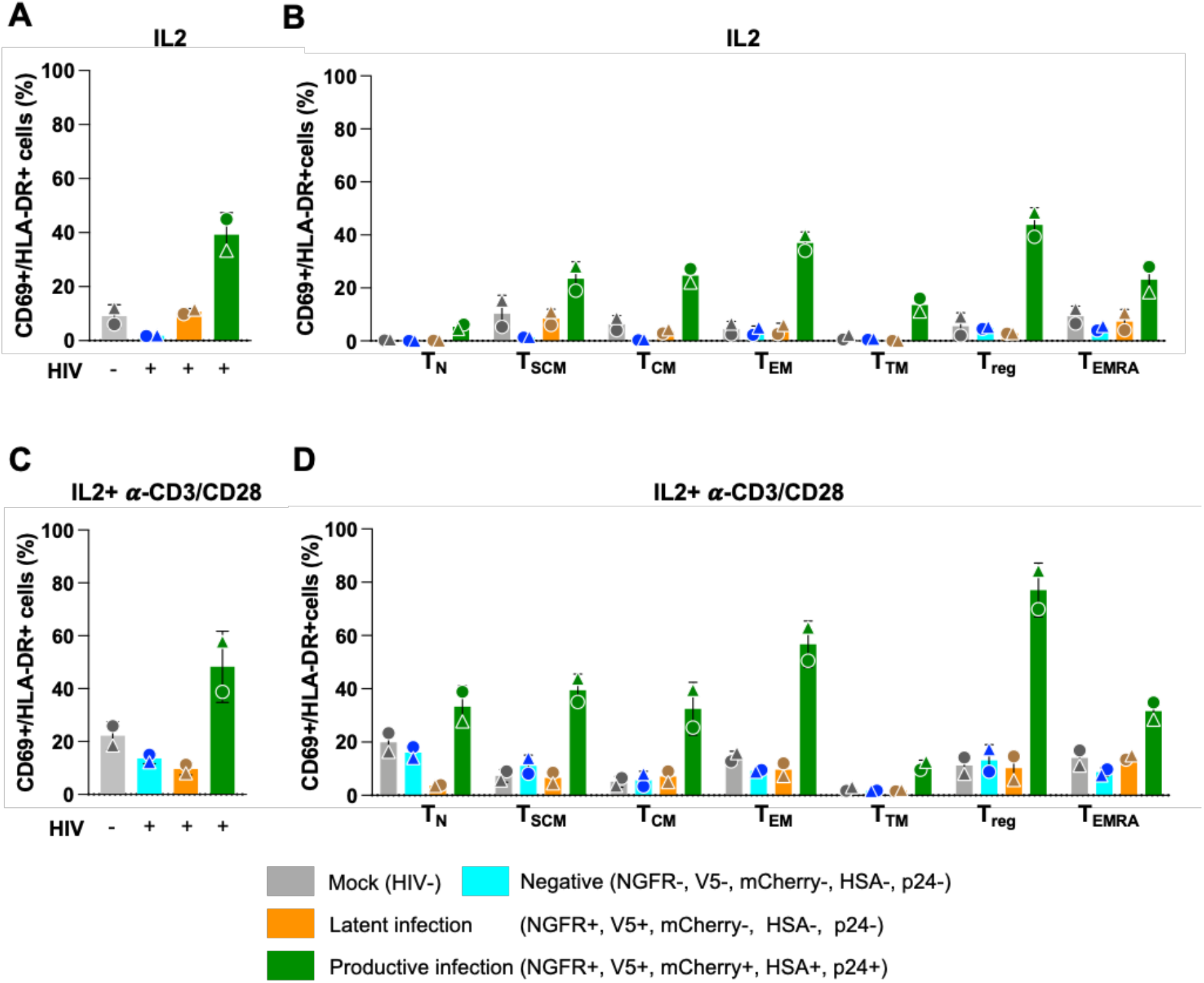
Validation of latent and productive infection by measuring early activation markers CD69 and HLA-DR in different CD4^+^ T cell subsets. (A) CD69+/HLA-DR+ expression on latently and productively pMorpheus-V5 infected primary human CD4^+^ T cells stimulated with IL-2 was determined using by CyTOF. Bar graphs represent levels of CD69 and HLA-DR in total CD4^+^ T cells. Each donor is indicated by a unique symbol. The infection status of each CD4^+^ T cell subset is color coded in the figure. Results are from two independent donors. (B) CD69+/HLA-DR+ expression on latently and productively pMorpheus-V5 infected CD4^+^ T subsets stimulated with IL-2 was determined. The levels of CD69+/HLA-DR+ in mock infected, reporter negative cells, latently infected and productively infected CD4^+^ T cell subsets stimulated were measured by CyTOF. The average of two individual donors is shown. (C) CD69+/HLA-DR+ expression on latently and productively pMorpheus-V5 infected primary human CD4^+^ T cells stimulated with IL-2 and *α*-CD3/CD28 beads was determined using by CyTOF. Bar graphs represent levels of CD69 and HLA-DR in total CD4^+^ T cells. Each donor is indicated by a unique symbol. The infection status of each CD4^+^ T cell subset is color coded in the figure. Results are from two independent donors. (D) CD69+/HLA-DR+ expression on latently and productively pMorpheus-V5 infected CD4^+^ T subsets stimulated with IL-2 and *α*-CD3/CD28 beads was determined. The levels of CD69+/HLA-DR+ in mock infected, reporter negative cells, latently infected and productively infected CD4^+^ T cell subsets were measured by CyTOF. The average of two individual donors is shown.

Taken together, our high-dimensional mass cytometry experiments detail the CD69/HLA-DR expression profiles of the CD4^+^ T cell subsets undergoing productive and latent infection. Furthermore, the stark differences in expression of the T cell activation markers CD69/HLA-DR represent an important independent confirmation of the ability of pMorpheus-V5 to distinguish productive from latent infections.

## Discussion

A more comprehensive understanding of the mechanisms supporting HIV latency is needed to guide HIV cure strategies. The ability to detect and enrich for latently infected primary human cells is essential to dissect mechanisms controlling HIV latency establishment, maintenance, and provirus reactivation. Dual-color HIV reporters are powerful tools to distinguish latently and productively infected cells from uninfected cells (Calvanese et al., 2013, Dahabieh et al., 2014, Battivelli et al., 2018). However, many of these dual-reporter HIV vectors proved to be suboptimal with respect to detecting HIV latently infected cells due to promoter interference between the viral LTR and the LTR independent promoter controlling expression of the phenotypic markers for productive and latent infection (Battivelli et al., 2018, Calvanese et al., 2013, Chavez et al., 2015, Dahabieh et al., 2013, Dahabieh et al., 2014, Hashemi et al., 2016, Kim et al., 2019, Matsuda et al., 2015). In this study, we built and characterized an improved version of a quadruple-reporter HIV (pMorpheus-V5) that efficiently identifies productive or latently infected primary human CD4^+^ T cells, stimulated with IL-2 and *α*-CD3/CD28 as well as IL-2 alone as a control (**Figure 1A, 1B**) with negligible promoter interference (**Figure 1C**, and **Figure S 2C**). Importantly, pMorpheus-V5 expresses full-length *Nef* as demonstrated by the efficient down-regulation of CD4^+^ exclusively in T cells harboring the LTR-active proviruses. Using this new system, we successfully enriched a population of latently infected primary CD4^+^ T cells without the need for fixation or intracellular staining. Infection with pMorpheus-V5 allows to distinguish cells harboring latent proviruses from those with productive viral replication using either by flow cytometry, and CyTOF (**Figures 2B, 2C, 2D, 4A, 4C, 4D**).

We noted that over 50% of the infected CD4^+^ T cells were latently infected in both conditions (**Figures 2C, 2D, 4C, 4D**), indicating that the level of latency establishment in primary CD4^+^ T cells might have been previously underestimated (Battivelli et al., 2018). This could be due to the fact that the two expression cassettes (LTR-dependent and independent) are positioned in a manner that minimizes promoter interference. In addition, the strength of the PGK promoter, which might overcome the action of epigenetic factors that cause HIV latency through LTR silencing could contribute to these observations. Moreover, our data indicate that HIV latency was established upon integration given that “early transcription” of multiply spliced RNA of regulatory proteins, *Tat* and *Rev*, is absent in latently infected cells (**Figure 3D**). Additionally, in both latent and productive infections, we show similar levels of integration (**Figure 3C**). By applying CyTOF high-dimensional immune profiling of primary human CD4^+^ T cells, we identified not only the CD4^+^ T cells subsets with high frequencies of latent infections but also the CD4^+^ populations most prone to latency establishment (**Figure 5A, 5B**). Finally, by measuring expression of activation markers CD69 and HLA-DR, we also present independent validation that V5-NGFR only positive cells are *bona fide* latently infected, which is a first important step towards improving our understanding of HIV latent reservoir formation.

T_EM_ and T_reg_ are the subsets where HIV latency establishment occurred with great efficiency (**Figure 5A, 5B**). Previous reports have shown that CD4^+^ T_EM_ cells contain the largest reservoir in which HIV reactivation has been observed (Kulpa et al., 2019, Buzon et al., 2014). CD4^+^ T_EM_ cells were also reported to harbor more intact proviruses (Hiener et al., 2017), while also maintaining a higher proliferation rate and a short half-life (Vukmanovic-Stejic et al., 2006, Farber et al., 2014). T_regs_ have also been reported to be latently infected representing an important HIV reservoir because of their potential to expand in chronically HIV-infected patients (Moreno-Fernandez et al., 2012). Moreover, T_regs_ were shown to contribute more to the HIV cellular reservoir than non-Tregs in HIV infected patients on ART (Tran et al., 2008). Their contribution to the rapid establishment of the HIV reservoir maybe due to their ability to reverse the immune activation status of CD4^+^ T cells (Moreno-Fernandez et al., 2012). Although with much lower frequency than for the T_CM_ and T_TM_ compartments, proviruses are almost always detected in naïve cells in individuals with controlled or uncontrolled HIV infection (Chomont et al., 2009, Bacchus et al., 2013, Douek et al., 2002, Lambotte et al., 2002, Zerbato et al., 2016). Similarly, we observed establishment of latency in the CD4^+^ T_N_ subset following either stimulation or activation (**Figure 5A, 5B, 5C**). Indeed, CD4^+^ T_N_ cells were the more prone subset to establish a latent infection. These results suggest that CD4^+^ T_N_ cells may serve as an important reservoir of LTR silent proviruses even though the overall frequency of latent infection is lower than that in the other CD4^+^ T cell subsets analyzed.

The molecular mechanism(s) supporting these differences require further targeted investigations, which now become feasible using pMorpheus-V5. In addition, we envision that pMorpheus-V5 will be an excellent tool for screening for drugs that selectively prevent or reverse latency establishment in primary human CD4^+^ T cell subpopulations.

## Materials And Methods

### Primary CD4^+^ T cell isolation and cell culture

Leukopaks were obtained from anonymous healthy blood donors from the New York Blood Center. Peripheral blood mononuclear cells were purified by Ficoll (Ficoll paque; GE Healthcare) density centrifugation and primary CD4^+^ T cells were purified by negative selection of CD4+ cells using magnetic beads (CD4+ T cell isolation kit, human; Miltenyi Biotec) according to the manufacturer’s instructions. CD4^+^ T cells were cultured in RPMI 1640 supplemented with 10% FBS (Gibco/Thermo Fisher), 100 IU/ml penicillin, 100 µg/mL streptomycin, 0.1 M HEPES, 2 mM L-glutamine in the presence of recombinant human IL-2 (20U/ml) (NIH AIDS Reagent Program, Division of AIDS, NIAID, NIH), or *α*-CD3/CD28 antibody-coated beads (Dynabeads Human T-Activator CD3/CD28; Thermo Fisher) in presence IL-2 (20U/ml) at 37°C in a 5% CO_2_ humidified incubator.

### Construction of pMorpheus-V5 and virus stock production

The genetic backbone of the pLAI2-V5-NGFR-HSA-mCherry-IRES-*Nef* (renamed in the text to pMorpheus-V5) is based on the pLAI2 HSA-mCherry-IRES-*Nef* full length viral construct (Young et al., 2018), which encodes the full-length HIV-1 strain LAI (Peden et al., 1991) as well as the HSA-mCherry-IRES-*Nef* cassette in the *Nef* ORF. The cassette encoding the PGK promoter followed by V5-NGFR (Wroblewska et al., 2018) was inserted in the envelope region encompassing the SalI and SmaI restriction sites. The construct was verified by sequencing. The virus stocks were produced by transfecting HEK 293T cells with 20 µg of pLAI2-V5-NGFR-HSA-mCherry-IRES-*Nef* and 5 µg of expression plasmid encoding the *Env* of pSV III-92 HT 594.1 subtype B virus R5 aid X4 (dual-tropic). After approximately 20 h, the transfection medium was changed, and the viral supernatants were collected at 48 and 72 hrs after transfection. The viral supernatants were filtered with 0.45 µm pore size and then concentrated using the Lenti-X concentrator (Takara) following manufacturer’s instruction.

The p24 Gag concentration of the pseudotyped viruses were determined using a commercial p24 ELISA (XpressBio) according to the manufacturers’ instructions. The optical density was read at 450 nm using a microtitration plate reader (VICTOR3_TM_, Perkin Elmer Precisely). Viral stocks were tested in triplicate. Aliquots of the pMorpheus-V5 pseudotyped with HIV *Env* stocks were stored at -80 °C until use.

### Infection of primary human CD4^+^ T cells with pMorpheus-V5

Primary human CD4^+^ T cells were incubated with 20U/ml interleukin-2 (IL-2) alone or in combination with *α*-CD3/CD28 antibody-coated beads (100 μL/30 x 10^6^ cells) (Gibco/Thermo Fisher) for 72 hours prior to infection with 100ng of p24 equivalent of pseudotyped pMorpheus-V5. For each donor, approximately ∼3 x 10^6^ cells of non-activated or activated cells were infected by spinoculation at 1,200 rpm for 2 hrs with equal amount of pMorpheus-V5 in presence of diethylaminoethyl (DEAE)-dextran (20 μg). After the spinoculation, the cells were washed and cultured in the presence of IL-2 (20U/ml) for five days. The number of infected cells were determined by quantifying NGFR, V5, mCherry, p24-expressing cells by Flow cytometry or by cytometry by time of flight (CyTOF).

As a negative control, we included in some experiments cells treated with FDA-approved Raltegravir (2μM, RAL, HIV Reagent Program, Manassas, VA, USA), which blocks HIV integration by preventing provirus integration into the host chromosome (Murray et al., 2007, Summa et al., 2008).

### Flow cytometry analysis

After infection with pMorpheus-V5, cells were pelleted, resuspended in PBS, and stained with the LIVE/DEAD Fixable Near-IR Dead Cell Stain Kit (Thermo Fisher Scientific) for 30 min at 4°C. Cells were washed and stained with antibodies against cell surface molecules in PBS containing 4% of human serum (Valley Biomedical). NGFR was detected with a 20 μl per million cells of Alexa Flour 647 conjugated *α*-NGFR (BD Bioscience), and V5 detected with a 1:100 dilution of fluorescein isothiocyanate (FITC)-conjugated *α*-mV5 (Invitrogen) in FACS buffer. Prior to flow cytometry analysis, cells were fixed in 1% (vol/vol) formaldehyde for 30 min at 4°C and analyzed on an LSRFortessa (BD Biosciences) flow cytometer. For intracellular p24 staining, cells were permeabilized for 30 min at 4°C with Perm/Wash buffer (BD Biosciences) and stained with *α*-p24 KC57 antibody (Beckman Coulter) for an additional 45 min at room temperature. Cells were washed and resuspended in PBS for subsequent flow cytometry analysis. CD4 downregulation was measured using PE-Cy5 antibody (Biolegend) with a 5 μl per million cells in 100 μl FACS buffer by flow cytometry. The viability of cells was first determined by the LIVE/DEAD Fixable Dead Cell stain (Thermo Fisher Scientific) with forward vs side scatter (FSC versus SSC). Data were gated on single CD3+ CD8-cells, and CD4 downregulation was assessed on cells expressing level of NGFR, V5, mCherry or p24. Gates were set by comparison with uninfected cells stained as above. An average of 10^5^ cells were acquired per sample, and data were analyzed using FlowJo software v10.7 (FlowJo, LLC).

### Measurements of HIV integration and transcription in FACS sorted CD4^+^ T cells

Viral stocks were pretreated with Dnase I (10U/ml, New England Biolabs Inc, MA, USA) and Benzonase nuclease (50U/ml, Millipore sigma) at room temperature for 1 hour before infection to remove background signal from plasmid DNA. Primary human CD4^+^ T cells stimulated with IL-2 alone or with *α*-CD3/CD28 antibody-coated beads (Gibco/Thermo Fisher) were infected for 5 days. Infected cells treated with 2μM of Raltegravir (AIDS research program) were used as negative control. Activated J-Lat 6.3 T cells were used as a positive control (Jordan et al., 2003). The J-Lat 6.3 cells were activated with phorbol 12-myristate 13-acetate (PMA, 16 nM, Sigma-Aldrich) and Ionomycin (0.5 μM, Sigma) (Spina et al., 2013).

pMorpheus-V5 infected cells were stained with LIVE/DEAD Fixable Near-IR Dead Cell Stain (Thermo Fisher Scientific) or V5-FITC Abs (Invitrogen) with a 1:100 dilution. Latently infected cells (V5_POS_ population), productively infected cells (mCherry_Pos_/V5_POS_ population), and negative cells (V5_neg_/mCherry_neg_ population) were sorted with a BD FACS Aria II Cell Sorter (BD Biosciences). After sorting the specific cell populations, cell pellets were lysed to isolate DNA and RNA using AllPrep DNA/RNA Kits (QIAGEN, Valencia, CA, USA) following the manufacturer’s protocol.

We measured HIV integration by *Alu*-1 PCR (Butler et al., 2001, Tan et al., 2006). Briefly, each PCR reaction contained 100 ng of DNA from each donor sample and 300 nM LM667 and 100 nM *Alu-*1 primers, and 40 μL Clone AMP to amplify integrated proviral DNA by thermocycler (BioRad). The PCR cycles initial denaturation at 95°C for 5 mins, then 20 cycles of 95°C for 10 secs, 55°C for 10 secs, and 72°C for 170 secs, and final extension of 72°C for 5 mins. We then diluted the amplicons from the first round of PCR 1:5. Each nested real-time PCR reaction included 2 μL of the diluted amplicons, and 300 nM each of primers LR1 and LR2, and 100 nM of ZXF-P probe containing Taqman universal master mix (PE-Applied Biosystems) and analyzed on a LightCycler (LightCycler 480 II, Roche Life Science, Roche Diagnostics Corporation, IN, USA). After incubation at 50°C for 2 min followed by 95°C for 10 min, 40 cycles of amplification were carried that included 15 sec at 95°C followed by 90 sec at 60°C. The 1μL *RNAseP* qPCR Probe reaction (Integrated DNA technologies), which included the 1μL DNA from each sample and Taqman universal master mix, was included as an internal control. The relative abundance of the integration events was calculated by delta-Ct method (*RnaseP*-sample) and shown on a log2 scale.

Multiply spliced HIV transcripts products were detected by semi-nested real-time quantitative-PCR as previously described (Pasternak et al., 2008, Zerbato et al., 2021). Briefly, we synthesized first-strand cDNA from cellular RNA isolated (100ng) from each sorted sample using iScript^TM^ cDNA synthesis kit (BioRad). Each rtPCR reaction included 25°C for 5 mins, 46°C for 20 mins, 95°C for 1min following the manufacturer’s instructions. Two rounds of PCR were performed with 2μL of cDNA, 10 μL of Clone AMP (Takara), and 300 nM each of primers ks1 and mf83. The PCR reactions included 15 cycles of 94°C for 3 mins, 55°C for 30 secs, 72°C for 1 min, and followed by final extension of 72°C for 5 mins. Subsequently, the semi-nested real-time q-PCR reactions (LightCycler 480 II, Roche Life Science, Roche Diagnostics Corporation, IN, USA) were performed with 1:5-diluted 1^st^ PCR product, and 300 nM of each primer mf83 and mf84, and 100 mM of the probe ks2-tq. Each reaction contained Taqman universal master mix (PE-Applied Biosystems). After incubation at 50°C for 2 min followed by 95°C for 10 min, 45 cycles of amplification were carried out including 15 sec at 95°C followed by 30 sec at 60°C. The *β*-actin reaction, which included the 2μL cDNA of each sample, 400 nM BGF, BGR primers, 300 nM BGX-P and Taqman universal master mix, was used as an internal control. Supplementary Table 1 summarizes the primers for the first PCR assay and subsequent nested qPCR assay. The quantity of multiply spliced HIV transcripts was determined by delta-Ct method (*β*-actin-sample) and shown on a log2 scale.

### Mass cytometry time-of-flight (CyTOF)

Ten million uninfected and infected human primary CD4^+^ T cells were stained with the Iridium (Ir) DNA intercalator, IdU_Ir193Di, to analyze DNA and cell cycle (Behbehani et al., 2012) and with the Rhodium (Rh) cationic nucleic acid intercalator, Rh103Di, to measure cellular viability. The cells were then washed in Cell Staining Media (CSM) buffer (0.5% BSA+0.02% NaN_3_ in PBS), stained with 2X CD298+ ß2M antibodies for barcoding (CD45_Cd111Di, CD45_Cd112Di, CD45_Cd114Di, CD45_Cd116Di, CD45_Pt195Di, CD45_Pt196Di), and pooled. The cells were washed with CSM buffer and resuspended in the membrane antibody mixture containing CD45RA_Nd143Di, CD45RO_Yb176Di, CD3_Er168DI, CD4_Nd145Di, CD24_Nd150Di, CD45_Y89Di, CD69_Nd144Di, CD127_Sm149Di, CD62L_Eu153Di, CD27_Gd155Di, CCR5_Gd156Di, CCR7_Gd156Di, CD25_Tm169Di, CD95_Yb171Di, CD28_Yb172Di, CXCR4_Yb173Di, Tim-3_Sm149Di, HLA-DR_Yb174Di, and PD-1_Sm147Di. After 30 min, the cells were washed in CSM. For fixation and permeabilization, the cells were fixed with paraformaldehyde in PBS for 10 min, followed by dropwise addition of cold methanol and storage at -80°C. The cells were washed in CSM containing heparin. Cells were resuspended in the phosphor staining antibody mixture containing Cyclin B_Dy163Di, pRb_Er166Di, pHistone3_LU175Di on ice for 30 min. The cells were washed in CSM and stained with intracellular antibody mixture containing FoxP3_Dy162Di, Ki-67_ Pr141Di, NGFR_Er170Di, V5_Sm151Di, mCherry_Nd140Di, HSA_Nd150Di, p24_KC57-RD_Ho165Di (BECKMAN COULTER) for 30 min on ice. The cells were washed and resuspended in Fix solution containing iridium intercalator. Between 200,000 and 550,000 CD4^+^ T cells were acquired per individual sample on the CyTOF2 or Helios mass cytometer (both Fluidigm). CyTOF settings were parameterized following the quality control of the instrument. After acquisition, the data were normalized using bead-based normalizing using CyTOF software (should list). The data was gated to exclude residual normalization beads, debris, and dead cells for clustering and high dimensional analysis. On acquisition, gating was done manually using the dot plot display on FlowJo or Cytobank. We gated only live cells for further analysis based on cellular viability (Rh103) and DNA content (Ir193). Seven different CD4^+^ T cell subpopulations were identified: Naïve (T_N_), Stem cell memory (T_SCM_), Central memory (T_CM_), Effector memory (T_EM_), Transitional memory (T_TM_), T Regulatory (T_REG_), and Effector memory T cells re-expressing CD45RA (T_EMRA_) cells **(Figure S3E)**. Figure S3B, S3C, S3D depicts the gating strategy for the CD45RO+ memory compartment. Briefly we first gated on the CD45RO cells followed by gating on CD27 and CCR7 to discriminate between T_CM_, T_EM_, and T_TM_ **(Figure S3B)** (Flynn et al., 2014). We distinguished T_N_ from T_SCM_ using the CD95 gate on CD45RA+, CCR7+, and CD27+ **(Figure S3C)** (Tabler et al., 2014, Klatt et al., 2014). T_REG_ were identified as being CD127-, Foxp3+ (Ndure et al., 2017), and T_EMRA_ (Tian et al., 2017) were identified by CCR7-, and CD45RA+ **(Figure S3D)** as previously described. The markers used to distinguish for each CD4^+^ T cell subset is in summarized **Figure S3E**.

### viSNE analysis

CyTOF data were debarcoded using Single Cell Debarcoder (Zunder et al., 2015) using post-assignment debarcode stringency filter and outlier trimming. We applied Cytobank viSNE (visualized t-stochastic neighbor embedding) (Amir el et al., 2013) and the Barnes-Hut implementation of the t-SNE algorithm to generate all viSNE plots of uninfected and pMorpheus-V5 infected cells from differently stimulated CD4^+^ T cells. The viSNE plots show the distribution of productively or latently infected cells present in each CD4^+^ T cell subpopulation based on an equal sampling of 66,000 cells from each file for cells stimulated with IL-2 alone or IL-2 and *α*-CD3/CD28 antibody-coated beads. In the viSNE plots, the position of each dot represents an individual cell. Dot-plot visualizations were performed using Cytobank software (Mountain View, CA).

### Statistical analysis

Statistical analyses were performed using GraphPad Prism 9 software. All results were presented as mean SD. Statistical analyses were performed using *t-test* or ANOVA for flow cytometry data. Differences with *P* values were indicated in the figures.

## ACKNOWLEDGEMENTS

We thank Dr. Adeeb Rahman and his team at the Human Immune Monitoring Core of Icahn School of Medicine at Mount Sinai for expertise and assistance of CyTOF. The following reagents were obtained through the NIH HIV Reagent Program, Division of AIDS, NIAID, NIH: J-Lat Full Length Cells (6.3), ARP-9846, contributed by Dr. Eric Verdin, and Raltegravir (Isentress; MK-0518), ARP-11680, contributed by Merck & Company, Inc. This research was supported in part by NIH/NIAID R01AI136916 (V.S.) and NIH AI150355 to L.C.F.M.

## Author Contributions

E.H.K. and V.S. designed research; E.H.K. performed all experiments; L.C.F.M. designed proviral vectors; E.H.K performed and analyzed data; E.H.K generated figures; E.H.K. and V.S. interpreted the data. B.D.B. provided critical reagents. M.S. and L.M. provided critical insight for flow data analysis and interpretation. E.H.K. drafted the manuscript. E.H.K, L.C.F.M., and V.S. revised the final manuscript. All authors approved the final version.

## Reagent Availability

Availability of the plasmid encoding the pMorpheus-V5 reporter construct is subject to Material Transfer Agreements.

## Supplementary Figures

**Figure S1.**
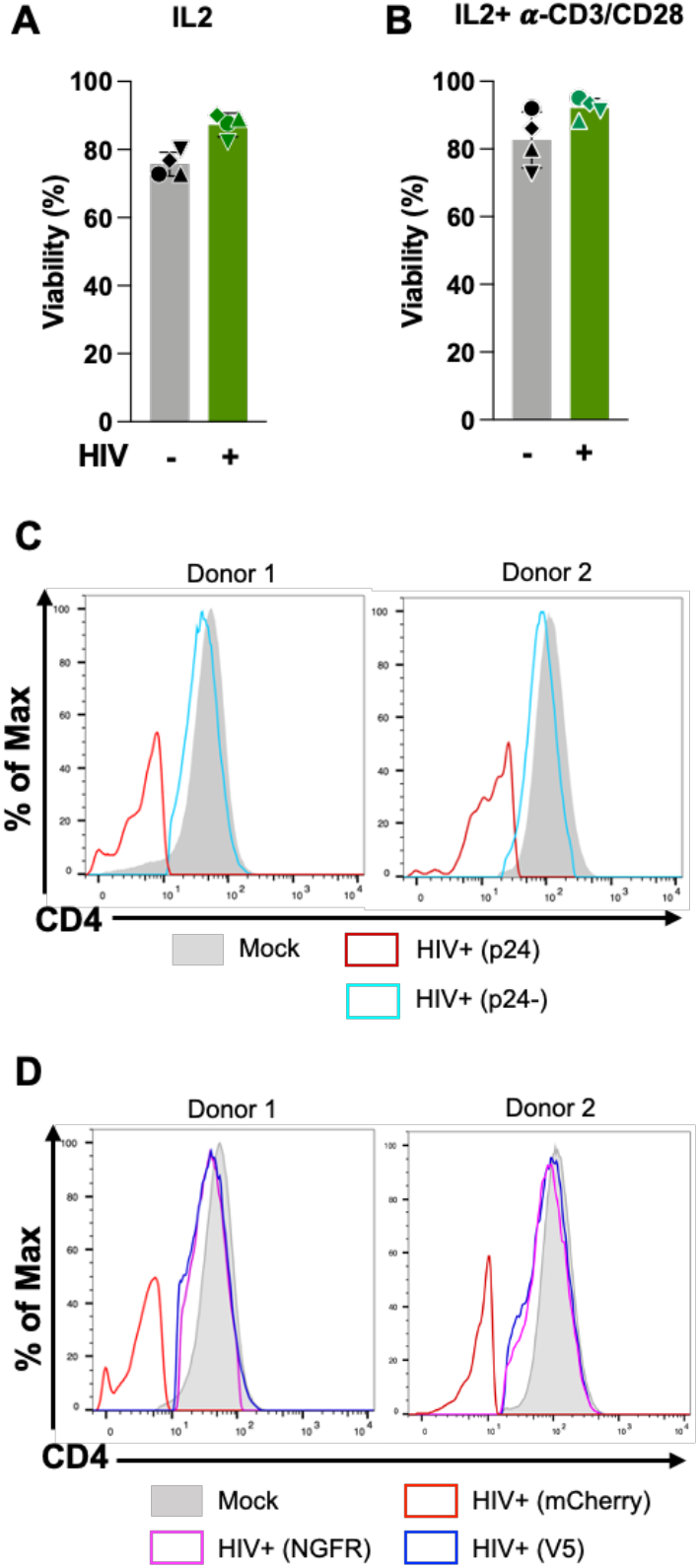
Validation of pMorpheus-V5 virus by flow cytometry. (A) CD4^+^ T cells from four donors were stimulated with IL-2 for 3 days and then infected with pMorpheus-V5. Five days after infection, cell viability was determined using the LIVE/DEAD Fixable Dead Cell stain by flow cytometry. The bar graphs show the percentage of live cells after infection. Error bars correspond to SD. \*\**p*=0.003. The average of four individual donors is shown (±SD). (B) CD4^+^ T cells from four donors were stimulated with IL-2 and *α*-CD3/CD28 for 3 days and then infected with pMorpheus-V5. Five days after infection, cell viability was determined by flow cytometry. The bar graphs show the percentage of live cells after infection. *p*=0.1. The average of four individual donors is shown (±SD). (C) CD4^+^ T cells from healthy donors were stimulated with IL-2 and *α*-CD3/CD28 beads and infected with pMorpheus-V5. Flow cytometry was performed 5 days post-infection. Histogram plots show the CD4 expression for uninfected cells (shaded light grey), p24_POS_ cells (red), or p24_NEG_ cells (light blue). The data were analyzed by FlowJo. Results are from four independent donors. (D) CD4^+^ T cells from healthy donors were stimulated with IL-2 and *α*-CD3/CD28 beads and infected with pMorpheus-V5. Flow cytometry was performed 5 days post-infection. CD4 expression levels of NGFR (magenta), V5 (blue), and mCherry (red) are shown as histogram plots. The data were analyzed by FlowJo. These results are representative of data from four independent donors.

**Figure S2.**
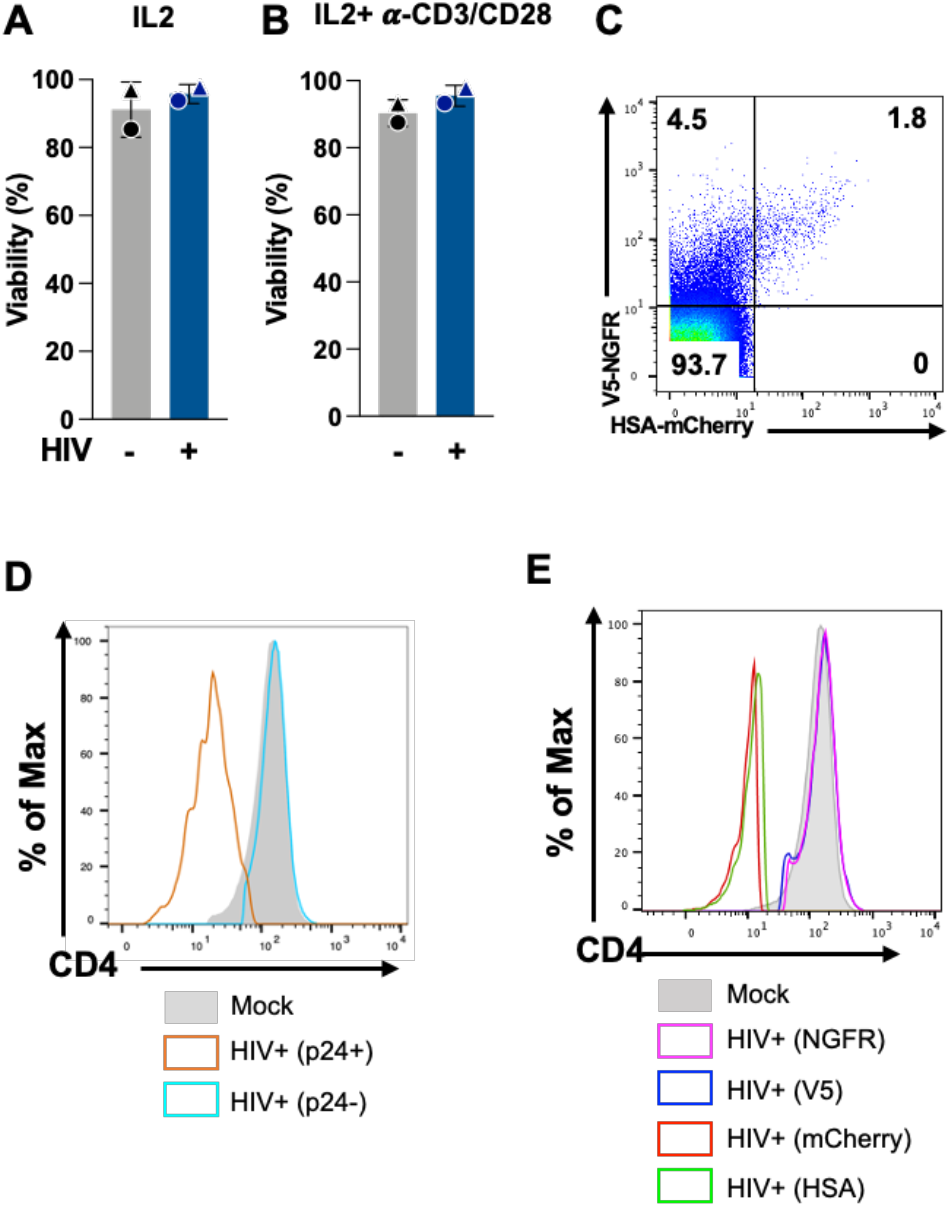
Validation of pMorpheus-V5 by CyTOF. (A) CD4^+^ T cells purified from PBMCs were cultured for 3 days in presence of IL-2 and then infected with pMorpheus-V5, which was pseudotyped with a dual-tropic HIV envelop. After 5 days of infection, cells were processed for CyTOF. pMorpheus-V5-infected primary human CD4^+^ T cells stimulated with IL-2 were analyzed by CyTOF to determine viability. The average of two individual donors is shown (±SD). Each donor is indicated by a unique symbol in the figure. (B) CD4^+^ T cells from two donors were stimulated with IL-2 and *α*-CD3/CD28 beads for 3 days and then infected with pMorpheus-V5. Cell viability was determined by CyTOF. Bar graphs represent percentage of live cells after infection. Error bars correspond to SD. The average of two individual donors is shown (±SD). Each donor is indicated by a unique symbol in the figure. (C) Cells were stimulated with IL-2 and *α*-CD3/CD28 antibody-coated beads prior to infection. Latently (V5_POS_, NGFR_NEG_) or productively (HSA_POS_, mCherry_Pos_+, V5_POS_, NGFR_POS_, p24_POS_) infected cells were identified by CyTOF five days after infection. These results are representative of data from two independent donors. (D) CD4^+^ T cells were purified from PBMCs and cultured for 3 days in presence of IL-2 and *α*-CD3/CD28 beads before infection with pMorpheus-V5. After 5 days of infection, the CD4 levels were quantified by CyTOF. Histogram plots show CD4 levels for uninfected cells (shaded light grey), p24_POS_ cells (orange), or p24_NEG_ cells (light blue) T cells. These results are representative of data from two independent donors. (E) IL-2 and *α*-CD3/CD28-stimulated CD4^+^ T cells were infected with pMorpheus-V5 for 5 days and analyzed by CyTOF to measure *Nef*-mediated downregulation of CD4 levels. NGFR (magenta), V5 (blue), and mCherry (red) cells were analyzed for CD4 levels. These results are representative of data from two independent donors.

**Figure S3.**
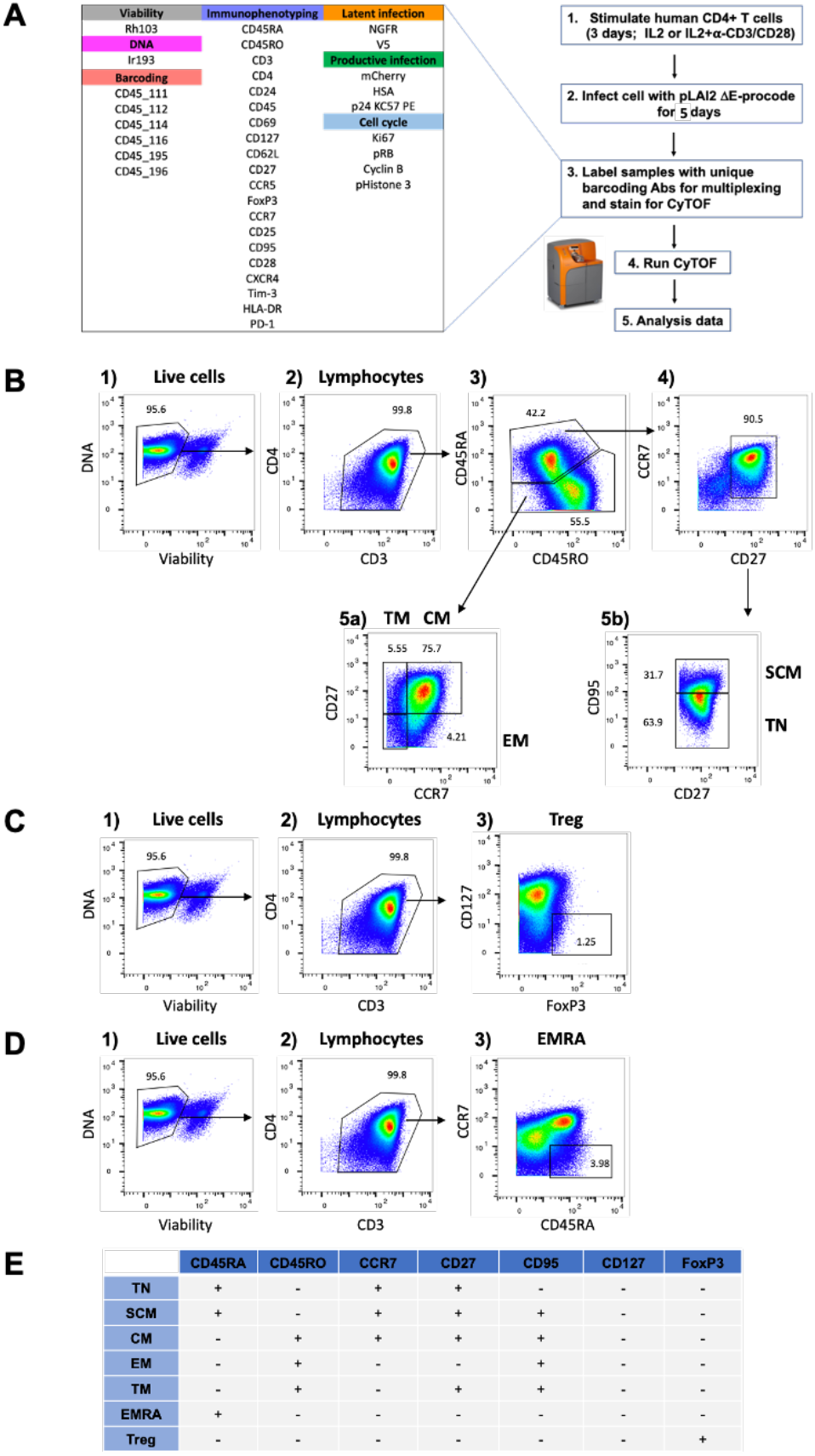
Overview of the experimental approach and gating strategy for CyTOF. (A) CD4^+^ T cells from two donors were stimulated with IL-2 or a combination of IL-2 with *α*-CD3/CD28 antibody-coated beads for 3 days and then infected with pMorpheus-V5. After 5 days of infection, cells were processed for CyTOF. Representative latent (NGFR_POS_, V5_POS_) or productive (HSA_POS_, mCherry_Pos_, NGFR_POS_, V5_POS_) infection profiles at 5 days post-infection for activated CD4^+^ T cells stimulated with IL-2. Individual donor cells were first barcoded using CD45 antibody and then mixed prior to staining with the panel of 29 phenotypic markers shown at left of the figure. (B) Outline of the gating strategy for analysis of the different CD4^+^ T cell populations. All live CD4^+^ T cell subsets were identified by DNA viability marker. T_CM_ are defined as CD3+. CD4+, CD45RA-, CCR7+, CD27+; T_EM_ are defined as CD3+, CD4+, CD45RA-, CCR7-, CD27-; T_TM_ are defined as CD3+, CD4+, CD45RA-, CCR7-, CD27+; T_N_ are defined as CD3+, CD4+, CD45RA+, CCR7+, CD27+, CD95-; TSCM are defined as CD3+, CD4+, CD45RA+, CCR7+ CD27+, CD95+. (C) T_reg_ are defined as CD3+, CD4+, CD127-, Foxp3+. (D) T_EMRA_ are defined as CD3+, CD4+, CCR7-, CD45RA+. (E) Table summarizing the phenotypic marker profile of each subset of CD4^+^ T cells examined.

**Figure S4.**
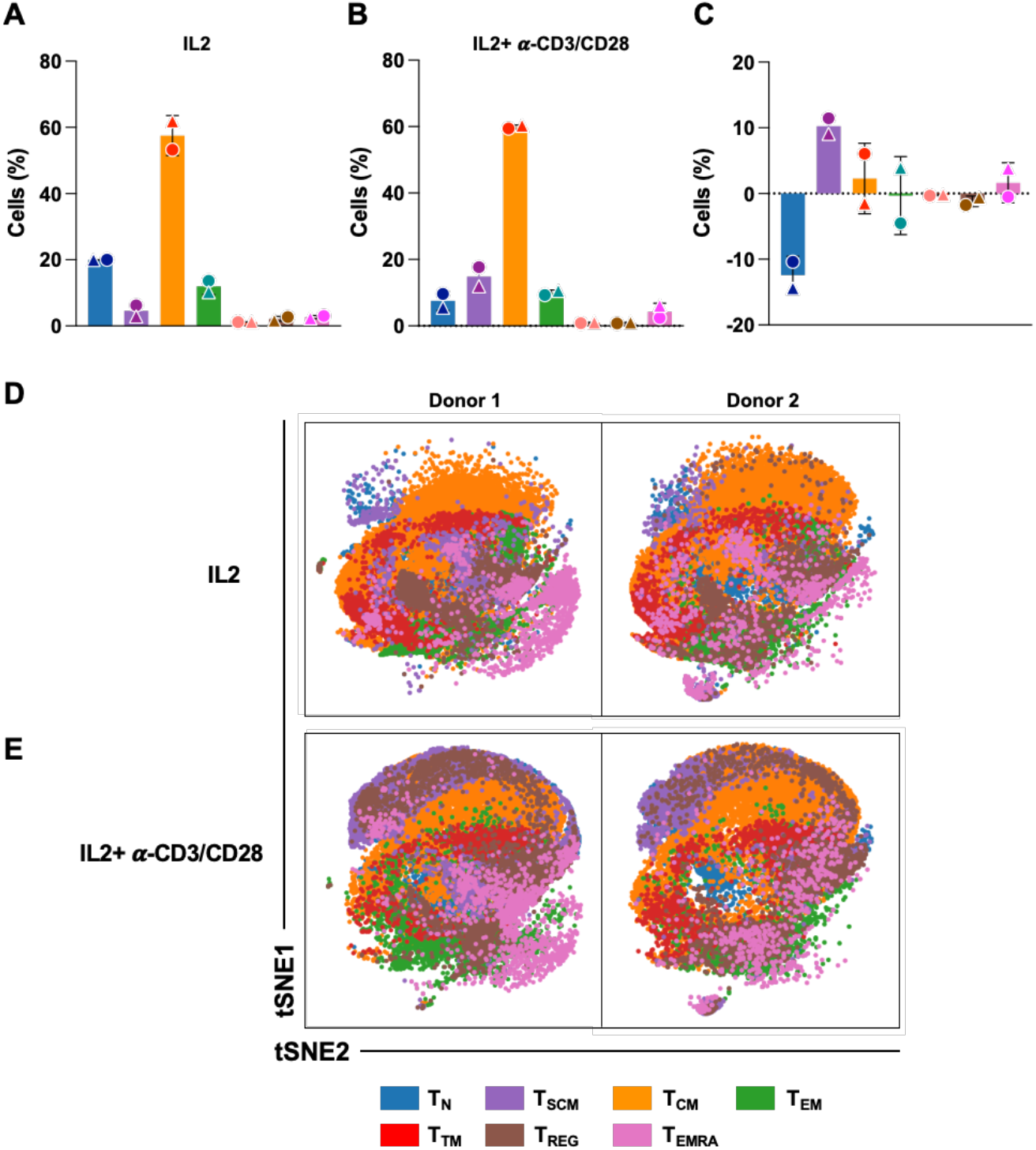
Distribution of CD4^+^ T cell subsets in presence of IL-2 stimulation or activation (in the absence of HIV infection) (A) CD4^+^ T cells from two donors were stimulated with IL-2 for 3 days. The bar graphs show the distribution of CD4^+^ T cell subsets from two different donors after 3 days of IL-2 stimulations as measured by CyTOF. These results are from two independent donors. Each donor is identified by a symbol. The legend shows the color code for each CD4^+^ T cell subset. The average of two individual donors is shown. (B) Primary human CD4^+^ T cells stimulated with IL-2 and *α*CD3/CD28 were anal by CyTOF. The distribution of CD4^+^ T cell subpopulations are shown as bar graphs. The results are from two independent donors. Each donor is indicated by a unique symbol. (C) Bar graph shows the fold change of CD4^+^ T cell subsets following IL-2 or IL-2 and *α*-CD3/CD28 bead stimulation as measured by CyTOF. The average of two individual donors is shown (±SD). Each donor is indicated by a unique symbol. (D) viSNE analysis showing absence of pMorpheus-V5 in CD4^+^ T cells in presence of IL-2 alone. The tSNE1 and tSNE2 axes are based on relevant markers of cell subsets defined by analysis with the Cytobank program. These results are from two independent donors. The legend shows the color code for each CD4^+^ T cell subset. (E) CD4^+^ T cells from two donors were stimulated with IL-2 and *α*-CD3/CD28 beads for three days. viSNE analysis arranged different CD4^+^ T cell subpopulations in presence of IL-2 and *α*-CD3/CD28 beads. The tSNE1 and tSNE2 axes are based on relevant markers of cell subsets defined by analysis with the Cytobank program. These results are from two independent donors. The legend shows the color code for each CD4^+^ T cell subset.

**Figure S5.**
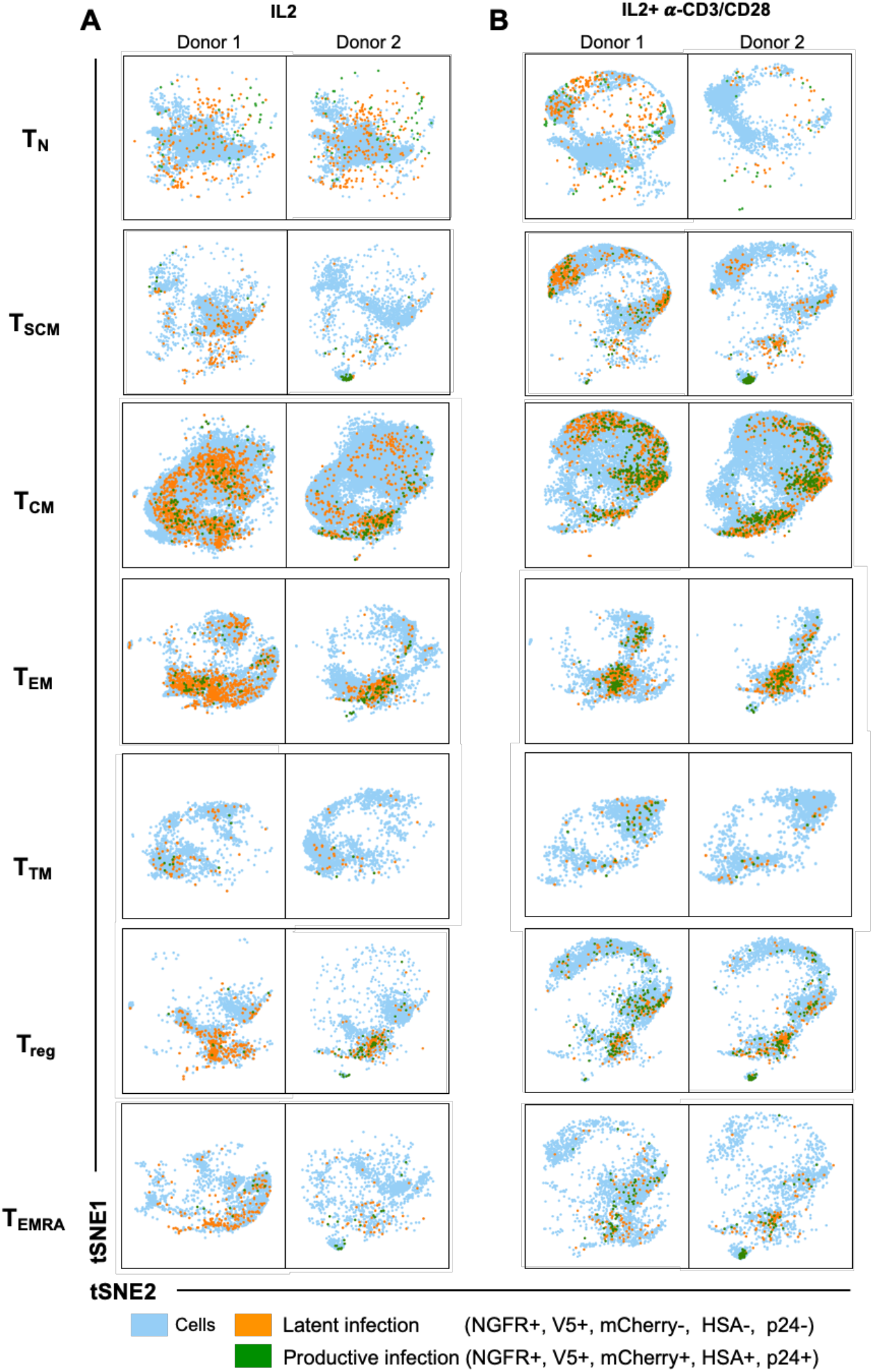
High dimensional CyTOF analysis reveals that distribution of CD4^+^ T cell subsets in presence of stimulations upon pMorpheus-V5 infection. (A) The infection of CD4^+^ T cell subsets from two different healthy donors after 3 days of IL-2 or IL-2 and *α*-CD3/CD28 bead stimulation followed by pMorpheus-V5 infection as measured by CyTOF. viSNE analysis arranged latently and productively infected CD4^+^ T cell subpopulations in presence of IL-2. These results are from two independent donors. Each infection is color coded in the figure. (B) CD4^+^ T cells from two healthy donors were stimulated with IL-2 and *α*-CD3/CD28 beads for three days and infected with pMorpheus-V5 for 5 days. The latent and productive infection in each subset was analyzed by CyTOF. viSNE analysis arranged latent and productive infection in different CD4^+^ T cell subpopulations following IL-2 and *α*-CD3/CD28 bead stimulation. The tSNE1 and tSNE2 axes are based on relevant markers of cell subsets defined by analysis with the Cytobank program. These results are from two independent donors. Each infection is color coded in the figure.

**Supplemental Table 1.**
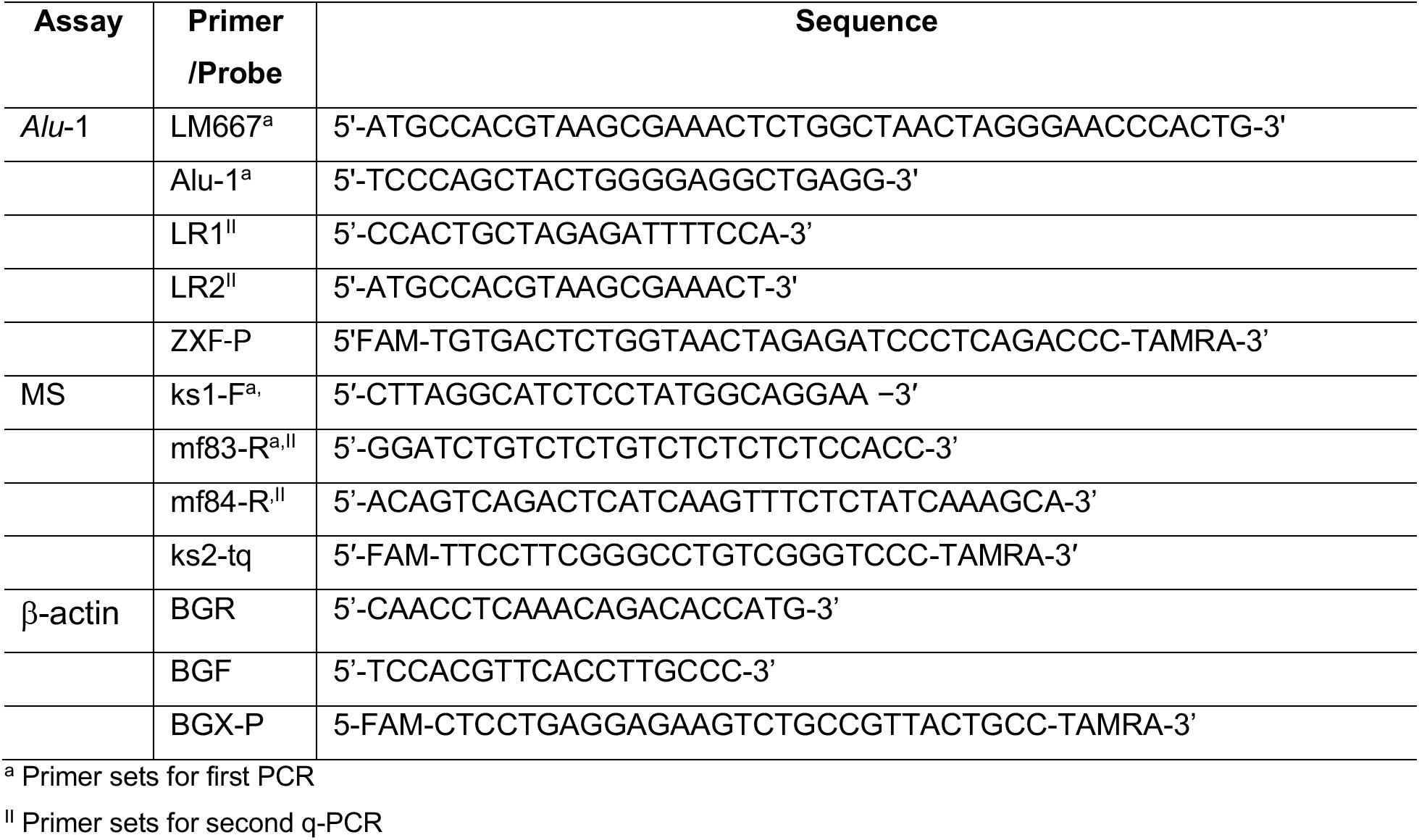
Oligonucleotide primers and probes used in this study.

## References

Amir El, A. D., Davis, K. L., Tadmor, M. D., Simonds, E. F., Levine, J. H., Bendall, S. C., Shenfeld, D. K., Krishnaswamy, S., Nolan, G. P. & Pe’er, D. 2013. viSNE enables visualization of high dimensional single-cell data and reveals phenotypic heterogeneity of leukemia. Nat Biotechnol, 31, 545–52.

Bacchus, C., Cheret, A., Avettand-Fenoel, V., Nembot, G., Melard, A., Blanc, C., Lascoux-Combe, C., Slama, L., Allegre, T., Allavena, C., Yazdanpanah, Y., Duvivier, C., Katlama, C., Goujard, C., Seksik, B.C., Leplatois, A., Molina, J. M., Meyer, L., Autran, B., Rouzioux, C. & GROUP, O. A. S. 2013. A single HIV-1 cluster and a skewed immune homeostasis drive the early spread of HIV among resting CD4+ cell subsets within one month post-infection. PLoS One, 8, e64219.

Battivelli, E., Dahabieh, M. S., Abdel-Mohsen, M., Svensson, J. P., Tojal Da Silva, I., Cohn, L. B., Gramatica, A., Deeks, S., Greene, W. C., Pillai, S. K. & Verdin, E. 2018. Distinct chromatin functional states correlate with HIV latency reactivation in infected primary CD4(+) T cells. Elife, 7.

Behbehani, G. K., Bendall, S. C., Clutter, M. R., Fantl, W. J. & Nolan, G. P. 2012. Single-cell mass cytometry adapted to measurements of the cell cycle. Cytometry A, 81, 552–66.

Butler, S. L., Hansen, M. S. & Bushman, F. D. 2001. A quantitative assay for HIV DNA integration in vivo. Nat Med, 7, 631–4.

Buzon, M. J., Sun, H., Li, C., Shaw, A., Seiss, K., Ouyang, Z., Martin-GAYO, E., Leng, J., Henrich, T. J., Li, J. Z., Pereyra, F., Zurakowski, R., Walker, B. D., Rosenberg, E. S., Yu, X. G. & Lichterfeld, M. 2014. HIV-1 persistence in CD4+ T cells with stem cell-like properties. Nat Med, 20, 139–42.

Cai, J., Gao, H., Zhao, J., Hu, S., Liang, X., Yang, Y., Dai, Z., Hong, Z. & Deng, K. 2021. Infection with a newly designed dual fluorescent reporter HIV-1 effectively identifies latently infected CD4(+) T cells. Elife, 10.

Calvanese, V., Chavez, L., Laurent, T., Ding, S. & Verdin, E. 2013. Dual-color HIV reporters trace a population of latently infected cells and enable their purification. Virology, 446, 283–92.

Cavrois, M., Banerjee, T., Mukherjee, G., Raman, N., Hussien, R., Rodriguez, B. A., Vasquez, J., Spitzer, M. H., Lazarus, N. H., Jones, J. J., Ochsenbauer, C., Mccune, J. M., Butcher, E. C., Arvin, A. M., Sen, N., Greene, W. C. & Roan, N. R. 2017. Mass Cytometric Analysis of HIV Entry, Replication, and Remodeling in Tissue CD4 + T Cells. Cell Rep, 20, 984–998.

Chavez, L., Calvanese, V. & Verdin, E. 2015. HIV Latency Is Established Directly and Early in Both Resting and Activated Primary CD4 T Cells. PLoS Pathog, 11, e1004955.

Chomont, N., El-Far, M., Ancuta, P., Trautmann, L., Procopio, F. A., Yassine-Diab, B., Boucher, G., Boulassel, M. R., Ghattas, G., Brenchley, J. M., Schacker, T. W., Hill, B. J., Douek, D. C., Routy, J. P., Haddad, E. K. & Sekaly, R. P. 2009. HIV reservoir size and persistence are driven by T cell survival and homeostatic proliferation. Nat Med, 15, 893–900.

Churchill, M. J., Deeks, S. G., Margolis, D. M., Siliciano, R. F. & Swanstrom, R. 2016. HIV reservoirs: what, where and how to target them. Nat Rev Microbiol, 14, 55–60.

Colin, L. & Van Lint, C. 2009. Molecular control of HIV-1 postintegration latency: implications for the development of new therapeutic strategies. Retrovirology, 6, 111.

Dahabieh, M. S., Ooms, M., Brumme, C., Taylor, J., Harrigan, P. R., Simon, V. & Sadowski, I. 2014. Direct non-productive HIV-1 infection in a T-cell line is driven by cellular activation state and NFkappaB. Retrovirology, 11, 17.

Dahabieh, M. S., Ooms, M., Simon, V. & Sadowski, I. 2013. A doubly fluorescent HIV-1 reporter shows that the majority of integrated HIV-1 is latent shortly after infection. J Virol, 87, 4716–27.

Douek, D. C., Brenchley, J. M., Betts, M. R., Ambrozak, D. R., Hill, B. J., Okamoto, Y., Casazza, J. P., Kuruppu, J., Kunstman, K., Wolinsky, S., Grossman, Z., Dybul, M., Oxenius, A., Price, D. A., Connors, M. & Koup, R. A. 2002. HIV preferentially infects HIV-specific CD4+ T cells. Nature, 417, 95–8.

Farber, D. L., Yudanin, N. A. & Restifo, N. P. 2014. Human memory T cells: generation, compartmentalization and homeostasis. Nat Rev Immunol, 14, 24–35.

Flynn, J. K., Paukovics, G., Cashin, K., Borm, K., Ellett, A., Roche, M., Jakobsen, M. R., Churchill, M. J. & Gorry, P.R. 2014. Quantifying susceptibility of CD4+ stem memory T-cells to infection by laboratory adapted and clinical HIV-1 strains. Viruses, 6, 709–26.

Gao, F., Morrison, S. G., Robertson, D. L., Thornton, C. L., Craig, S., Karlsson, G., Sodroski, J., Morgado, M., Galvao-Castro, B., Von Briesen, H., Beddows, S., Weber, J., Sharp, P. M., Shaw, G. M. & Hllahn, B. H. 1996. Molecular cloning and analysis of functional envelope genes from human immunodeficiency virus type 1 sequence subtypes A through G. The WHO and NIAID Networks for HIV Isolation and Characterization. J Virol, 70, 1651–67.

Garcia, J. V. & Miller, A. D. 1991. Serine phosphorylation-independent downregulation of cell-surface CD4 by nef. Nature, 350, 508–11.

Grau-Exposito, J., Luque-Ballesteros, L., Navarro, J., Curran, A., Burgos, J., Ribera, E., Torrella, A., Planas, B., Badia, R., Martin-Castillo, M., Fernandez-Sojo, J., Genesca, M., Falco, V. & Buzon, M. J. 2019. Latency reversal agents affect differently the latent reservoir present in distinct CD4+ T subpopulations. PLoS Pathog, 15, e1007991.

Guy, B., Kieny, M. P., Riviere, Y., Le PEUCH, C., Dott, K., Girard, M., Montagnier, L. & Lecocq, J. P. 1987. HIV F/3’ orf encodes a phosphorylated GTP-binding protein resembling an oncogene product. Nature, 330, 266–9.

Hashemi, F. B., Barreto, K., Bernhard, W., Hashemi, P., Lomness, A. & Sadowski, 2016. HIV Provirus Stably Reproduces Parental Latent and Induced Transcription Phenotypes Regardless of the Chromosomal Integration Site. J Virol, 90, 5302–14.

Hiener, B., Horsburgh, B. A., Eden, J. S., Barton, K., Schlub, T. E., Lee, E., Von Stockenstrom, S., Odevall, L., Milush, J. M., Liegler, T., Sinclair, E., Hoh, R., Boritz, E. A., Douek, D., Fromentin, R., Chomont, N., Deeks, S. G., Hecht, F.M. & Palmer, S. 2017. Identification of Genetically Intact HIV-1 Proviruses in Specific CD4(+) T Cells from Effectively Treated Participants. Cell Rep, 21, 813–822.

Jefferys, S. R., Burgos, S. D., Peterson, J. J., Selitsky, S. R., Turner, A. W., James, L. I., Tsai, Y. H., Coffey, A. R., Margolis, D. M., Parker, J. & Browne, E. P. 2021. Epigenomic characterization of latent HIV infection identifies latency regulating transcription factors. PLoS Pathog, 17, e1009346.

Jones, S., Peng, P. D., Yang, S., Hsu, C., Cohen, C. J., Zhao, Y., Abad, J., Zheng, Z., Rosenberg, S. A. & Morgan, R.A. 2009. Lentiviral vector design for optimal T cell receptor gene expression in the transduction of peripheral blood lymphocytes and tumor-infiltrating lymphocytes. Hum Gene Ther, 20, 630–40.

Jordan, A., Bisgrove, D. & Verdin, E. 2003. HIV reproducibly establishes a latent infection after acute infection of T cells in vitro. Embo J, 22, 1868–77.

Kim, Y., Cameron, P. U., Lewin, S. R. & Anderson, J. L. 2019. Limitations of dual-fluorescent HIV reporter viruses in a model of pre-activation latency. J Int AIDS Soc, 22, e25425.

Klatt, N. R., Bosinger, S. E., Peck, M., Richert-Spuhler, L. E., Heigele, A., Gile, P., Patel, N., Taaffe, J., Julg, B., Camerini, D., Torti, C., Martin, J. N., Deeks, S. G., Sinclair, E., Hecht, F. M., Lederman, M. M., Paiardini, M., Kirchhoff, F., Brenchley, J. M., Hunt, P. W. & Silvestri, G. 2014. Limited HIV infection of central memory and stem cell memory CD4+ T cells is associated with lack of progression in viremic individuals. PLoS Pathog, 10, e1004345.

Kotecha, N., Krutzik, P. O. & Irish, J. M. 2010. Web-based analysis and publication of flow cytometry experiments. Curr Protoc Cytom, Chapter 10, Unit10 17.

Kulpa, D. A., Talla, A., Brehm, J. H., Ribeiro, S. P., Yuan, S., Bebin-Blackwell, A. G., Miller, M., Barnard, R., Deeks, S. G., Hazuda, D., Chomont, N. & Sekaly, R.P. 2019. Differentiation into an Effector Memory Phenotype Potentiates HIV-1 Latency Reversal in CD4(+) T Cells. J Virol, 93.

Lambotte, O., Demoustier, A., De Goer, M. G., Wallon, C., Gasnault, J., Goujard, C., Delfraissy, J. F. & Taoufik, Y. 2002. Persistence of replication-competent HIV in both memory and naive CD4 T cell subsets in patients on prolonged and effective HAART. AIDS, 16, 2151–7.

Marini, A., Harper, J. M. & Romerio, F. 2008. An in vitro system to model the establishment and reactivation of HIV-1 latency. J Immunol, 181, 7713–20.

Matsuda, Y., Kobayashi-ISHIHARA, M., Fujikawa, D., Ishida, T., Watanabe, T. & Yamagishi, M. 2015. Epigenetic heterogeneity in HIV-1 latency establishment. Sci Rep, 5, 7701.

Mbonye, U. & Karn, J. 2014. Transcriptional control of HIV latency: cellular signaling pathways, epigenetics, happenstance and the hope for a cure. Virology, 454-455, 328–39.

Moreno-Fernandez, M. E., Presicce, P. & Chougnet, C. A. 2012. Homeostasis and function of regulatory T cells in HIV/SIV infection. J Virol, 86, 10262–9.

Moso, M. A., Anderson, J. L., Adikari, S., Gray, L. R., Khoury, G., Chang, J. J., Jacobson, J. C., Ellett, A. M., Cheng, W. J., Saleh, S., Zaunders, J. J., Purcell, D. F. J., Cameron, P. U., Churchill, M. J., Lewin, S. R. & Lu, H. K. 2019. HIV latency can be established in proliferating and nonproliferating resting CD4+ T cells in vitro: implications for latency reversal. AIDS, 33, 199–209.

Murray, J. M., Emery, S., Kelleher, A. D., Law, M., Chen, J., Hazuda, D. J., Nguyen, B. Y., Teppler, H. & Cooper, D. A. 2007. Antiretroviral therapy with the integrase inhibitor raltegravir alters decay kinetics of HIV, significantly reducing the second phase. Aids, 21, 2315–21.

Ndure, J., Noho-Konteh, F., Adetifa, J. U., Cox, M., Barker, F., Le, M. T., Sanyang, L. C., Drammeh, A., Whittle, H. C., Clarke, E., Plebanski, M., Rowland-Jones, S. L. & Flanagan, K.L. 2017. Negative Correlation between Circulating CD4(+)FOXP3(+)CD127(-) Regulatory T Cells and Subsequent Antibody Responses to Infant Measles Vaccine but Not Diphtheria-Tetanus-Pertussis Vaccine Implies a Regulatory Role. Front Immunol, 8, 921.

Pasternak, A. O., Adema, K. W., Bakker, M., Jurriaans, S., Berkhout, B., Cornelissen, M. & Lukashov, V. V. 2008. Highly sensitive methods based on seminested real-time reverse transcription-PCR for quantitation of human immunodeficiency virus type 1 unspliced and multiply spliced RNA and proviral DNA. J Clin Microbiol, 46, 2206–11.

Pearson, R., Kim, Y. K., Hokello, J., Lassen, K., Friedman, J., Tyagi, M. & Karn, J. 2008. Epigenetic silencing of human immunodeficiency virus (HIV) transcription by formation of restrictive chromatin structures at the viral long terminal repeat drives the progressive entry of HIV into latency. J Virol, 82, 12291–303.

Peden, K., Emerman, M. & Montagnier, L. 1991. Changes in growth properties on passage in tissue culture of viruses derived from infectious molecular clones of HIV-1LAI, HIV-1MAL, and HIV-1ELI. Virology, 185, 661–72.

Sahu, G. K., Lee, K., Ji, J., Braciale, V., Baron, S. & Cloyd, M. W. 2006. A novel in vitro system to generate and study latently HIV-infected long-lived normal CD4+ T-lymphocytes. Virology, 355, 127–37.

Siliciano, R. F. & Greene, W. C. 2011. HIV latency. Cold Spring Harb Perspect Med, 1, a007096.

Spina, C. A., Anderson, J., Archin, N. M., Bosque, A., Chan, J., Famiglietti, M., Greene, W. C., Kashuba, A., Lewin, S. R., Margolis, D. M., Mau, M., Ruelas, D., Saleh, S., Shirakawa, K., Siliciano, R. F., Singhania, A., Soto, P. C., Terry, V. H., Verdin, E., Woelk, C., Wooden, S., Xing, S. & Planelles, V. 2013. An in-depth comparison of latent HIV-1 reactivation in multiple cell model systems and resting CD4+ T cells from aviremic patients. PLoS Pathog, 9, e1003834.

Summa, V., Petrocchi, A., Bonelli, F., Crescenzi, B., Donghi, M., Ferrara, M., Fiore, F., Gardelli, C., Gonzalez PAZ, O., Hazuda, D. J., Jones, P., Kinzel, O., Laufer, R., Monteagudo, E., Muraglia, E., Nizi, E., Orvieto, F., Pace, P., Pescatore, G., Scarpelli, R., Stillmock, K., Witmer, M. V. & Rowley, M. 2008. Discovery of raltegravir, a potent, selective orally bioavailable HIV-integrase inhibitor for the treatment of HIV-AIDS infection. J Med Chem, 51, 5843–55.

Tabler, C. O., Lucera, M. B., Haqqani, A. A., Mcdonald, D. J., Migueles, S. A., Connors, M. & Tilton, J. C. 2014. CD4+ memory stem cells are infected by HIV-1 in a manner regulated in part by SAMHD1 expression. J Virol, 88, 4976–86.

Tan, W., Dong, Z., Wilkinson, T. A., Barbas, C. F., 3RD & Chow, S. A. 2006. Human immunodeficiency virus type 1 incorporated with fusion proteins consisting of integrase and the designed polydactyl zinc finger protein E2C can bias integration of viral DNA into a predetermined chromosomal region in human cells. J Virol, 80, 1939–48.

Tian, Y., Babor, M., Lane, J., Schulten, V., Patil, V. S., Seumois, G., Rosales, S. L., Fu, Z., Picarda, G., Burel, J., Zapardiel-Gonzalo, J., Tennekoon, R. N., De Silva, A. D., Premawansa, S., Premawansa, G., Wijewickrama, A., Greenbaum, J. A., Vijayanand, P., Weiskopf, D., Sette, A. & Peters, B. 2017. Unique phenotypes and clonal expansions of human CD4 effector memory T cells re-expressing CD45RA. Nat Commun, 8, 1473.

Tran, T. A., De Goer De Herve, M. G., Hendel-Chavez, H., Dembele, B., Le Nevot, E., Abbed, K., Pallier, C., Goujard, C., Gasnault, J., Delfraissy, J. F., Balazuc, A. M. & Taoufik, Y. 2008. Resting regulatory CD4 T cells: a site of HIV persistence in patients on long-term effective antiretroviral therapy. PLoS One, 3, e3305.

Vukmanovic-STEJIC, M., Zhang, Y., Cook, J. E., Fletcher, J. M., Mcquaid, A., Masters, J. E., Rustin, M. H., Taams, L. S., Beverley, P. C., Macallan, D. C. & Akbar, A. N. 2006. Human CD4+ CD25hi Foxp3+ regulatory T cells are derived by rapid turnover of memory populations in vivo. J Clin Invest, 116, 2423–33.

Weinberger, L. S., Dar, R. D. & Simpson, M. L. 2008. Transient-mediated fate determination in a transcriptional circuit of HIV. Nat Genet, 40, 466–70.

Wroblewska, A., Dhainaut, M., Ben-Zvi, B., Rose, S. A., Park, E. S., Amir, E. D., Bektesevic, A., Baccarini, A., Merad, M., Rahman, A. H. & Brown, B. D. 2018. Protein Barcodes Enable High-Dimensional Single-Cell CRISPR Screens. Cell, 175, 1141–1155 e16.

Young, G. R., Terry, S. N., Manganaro, L., Cuesta-Dominguez, A., Deikus, G., Bernal-Rubio, D., Campisi, L., Fernandez-Sesma, A., Sebra, R., Simon, V. & Mulder, L. C. F. 2018. HIV-1 Infection of Primary CD4(+) T Cells Regulates the Expression of Specific Human Endogenous Retrovirus HERV-K (HML-2) Elements. J Virol, 92.

Zerbato, J. M., Khoury, G., Zhao, W., Gartner, M. J., Pascoe, R. D., Rhodes, A., Dantanarayana, A., Gooey, M., Anderson, J., Bacchetti, P., Deeks, S. G., Mcmahon, J., Roche, M., Rasmussen, T. A., Purcell, D. F. & Lewin, S. R. 2021. Multiply spliced HIV RNA is a predictive measure of virus production ex vivo and in vivo following reversal of HIV latency. EBioMedicine, 65, 103241.

Zerbato, J. M., Serrao, E., Lenzi, G., Kim, B., Ambrose, Z., Watkins, S. C., Engelman, A. N. & Sluis-CREMER, N. 2016. Establishment and Reversal of HIV-1 Latency in Naive and Central Memory CD4+ T Cells In Vitro. J Virol, 90, 8059–73.

Zunder, E. R., Finck, R., Behbehani, G. K., Amir El, A. D., Krishnaswamy, S., Gonzalez, V. D., Lorang, C. G., Bjornson, Z., Spitzer, M. H., Bodenmiller, B., Fantl, W. J., Pe’ER, D. & Nolan, G. P. 2015. Palladium-based mass tag cell barcoding with a doublet-filtering scheme and single-cell deconvolution algorithm. Nat Protoc, 10, 316–33.

